# Aquila: diploid personal genome assembly and comprehensive variant detection based on linked reads

**DOI:** 10.1101/660605

**Authors:** Xin Zhou, Lu Zhang, Ziming Weng, David L. Dill, Arend Sidow

**Affiliations:** Department of Computer Science, Stanford University; Department of Pathology, Stanford University; Department of Genetics, Stanford University

## Abstract

Variant discovery in personal, whole genome sequence data is critical for uncovering the genetic contributions to health and disease. We introduce a new approach, Aquila, that uses linked-read data for generating a high quality diploid genome assembly, from which it then comprehensively detects and phases personal genetic variation. Assemblies cover >95% of the human reference genome, with over 98% in a diploid state. Thus, the assemblies support detection and accurate genotyping of the most prevalent types of human genetic variation, including single nucleotide polymorphisms (SNPs), small insertions and deletions (small indels), and structural variants (SVs), in all but the most difficult regions. All heterozygous variants are phased in blocks that can approach arm-level length. The final output of Aquila is a diploid and phased personal genome sequence, and a phased VCF file that also contains homozygous and a few unphased heterozygous variants. Aquila represents a cost-effective evolution of whole-genome reconstruction that can be applied to cohorts for variation discovery or association studies, or to single individuals with rare phenotypes that could be caused by SVs or compound heterozygosity.

## Introduction

Despite recent advances, quantifying the contribution of genetic variation to specific disease risk is a stubborn biomedical problem that remains far from solved. In general, understanding the relationship between genotype and phenotype requires complete ascertainment of genotype, which for humans has yet to be achieved in a scalable fashion. At this stage in technology development, DNA sequencing still faces a vexing tradeoff between cost and completeness so that discovery of variation in larger cohorts is limited to SNPs and small indels. In fact, the relatively low cost of Illumina-based short-fragment whole genome sequencing and the even lower cost of exomes and genotyping arrays has caused considerable ascertainment bias such that the vast majority of genotype-phenotype associations focus on SNPs with small effect, even though the undetected larger variation is known to involve roughly as many bases in our genomes as SNPs and is therefore predicted to have significant phenotypic impact as well (Redon et al. 2006; Weischenfeldt et al. 2013). Also generally missing is the phasing of genetic variation, which is similarly important for estimation of phenotypic impact, as the distinction between cis and trans compound heterozygotes in an essential locus can mean the difference between health and disease (Lupski et al. 2010) and is likely to modulate risk of multigenic disease as well (Jiang et al. 2018).

Single-molecule sequencing approaches, particularly Pacific Biosciences (PacBio) and Oxford Nanopore Technologies (ONT), provide potential solutions, as long-range information allows accurate detection of SVs and phasing (Coster et al. 2018; Huddleston et al. 2017; Cretu Stancu et al. 2017). However, the drawback of both approaches is that they exhibit poor base-pair level accuracy, leading to high error rates for SNPs and imprecise breakpoint estimation for small indels and SVs. A widely applied solution has been to supplement long reads with higher quality short read data, but these ensemble approaches are difficult to scale to larger cohorts due to the complexity of data generation, integration, and analysis, and have therefore been limited to small sample sizes in proof-of-principle studies (Rhoads and Au, 2015; Fan et al., 2017). A solution to making long reads more accurate is to sequence the same single molecule multiple times to reduce error, for example as implemented in the PacBio circular consensus sequencing (CCS) approach (Travers et al. 2010; Larsen et al. 2014; Wenger et al. 2019). However, CCS requires several-fold oversampling of the same molecule, a currently expensive proposition for anything but small sample sizes.

A relatively recent addition to the DNA sequencing ecosystem has been pioneered by 10X Genomics, wherein the original large molecules of a gentle DNA preparation are partitioned into microfluidic compartments (Zheng et al. 2016; Wang et al. 2019). Via a series of within-compartment molecular biology and subsequent standard steps of library construction and sequencing, barcoded short reads are produced that retain the long-range information of the long fragments of the initial DNA extract. Due to the combination of high base pair-level sequence accuracy and long-range information, 10x/Illumina data therefore support excellent SNP and small indel detection and phasing (Zheng et al. 2016), as well as breakpoint detection of large events in cancer (Zheng et al. 2016; Spies et al. 2017; Elyanow et al. 2017). For diploid genome reconstruction, 10x developed the *de novo* assembler, Supernova, which has been shown to produce whole human genome assemblies from 56-fold coverage 10x/Illumina data (Weisenfeld et al. 2017; Zhang et al. 2019a).

The application of assembly approaches to human genomes has been limited even though they allow powerful identification of SVs (Nattestad and Schatz, 2016, n.d.; Fan et al., 2017; Wala et al., 2018). Long-read based assemblies, such as those from PacBio data performed by FALCON-Unzip (Chin et al. 2016), exhibit respectable contiguity and variant detection but still suffer from high cost (Rhoads and Au 2015). Supernova assemblies based on 10x/Illumina data are less expensive and allow detection of all types of variation but power is limited because a substantial fraction of the genome is not assembled in a diploid state and genotyping error is still high (Zhang et al. 2019b). Overall, cost-effective assembly-based approaches still suffer from incomplete resolution of the diploid genome and limited power of variant detection in a personal genome. On the other hand, assembly-based approaches have two advantages: detection of variants is greatly simplified to pairwise alignments rather than complicated read-map based inference, which is particularly challenging for indels; and the detection of sequences not present in the reference.

Compared to reference-based approaches (Pop 2004), the competitive disadvantage of *de novo* assembly methods is that they disregard the high information content of the reference. Depending on genetic background, greater than 99.5% of anybody’s two haplomes is identical to the reference, which therefore constitutes a highly accurate scaffold for personal genomes. It stands to reason that, in principle, an assembly-based method that incorporates information from the reference sequence should combine the advantages of both approaches. We were therefore motivated to develop a new approach to accomplish these three goals via a reference-assisted, assembly-based approach: high quality diploid personal genome reconstruction; accurate detection of SNPs, indels and SVs; phasing of all types of variants.

Our method, Aquila, makes use of the reference genome by performing local assembly in small chunks separately for each haplotype, yielding a diploid whole genome consisting of local, phased, contigs whose scaffolding is provided by the reference sequence. It then discovers the most important types of variation on the basis of pairwise alignment to the reference, and infers phasing for all types of assembled variants through previous long-range phasing information. We tested its performance with six libraries of 10x linked-reads data for NA12878 and NA24385 individuals. It offers excellent small indel and SV detection at virtually no compromise for SNP detection, as well as highly accurate phasing of the vast majority of heterozygous variants, at reasonable reagent and computational costs.

## Results

### Aquila architecture and italic

Aquila consists of four stages (Figure 1A): Haplotyping and sorting inferred long fragments and their reads into the two parental bins, locally partitioning reads of each bin, assembling each local partition into sequence contigs, and finally variant calling and phasing.

**Figure 1.**
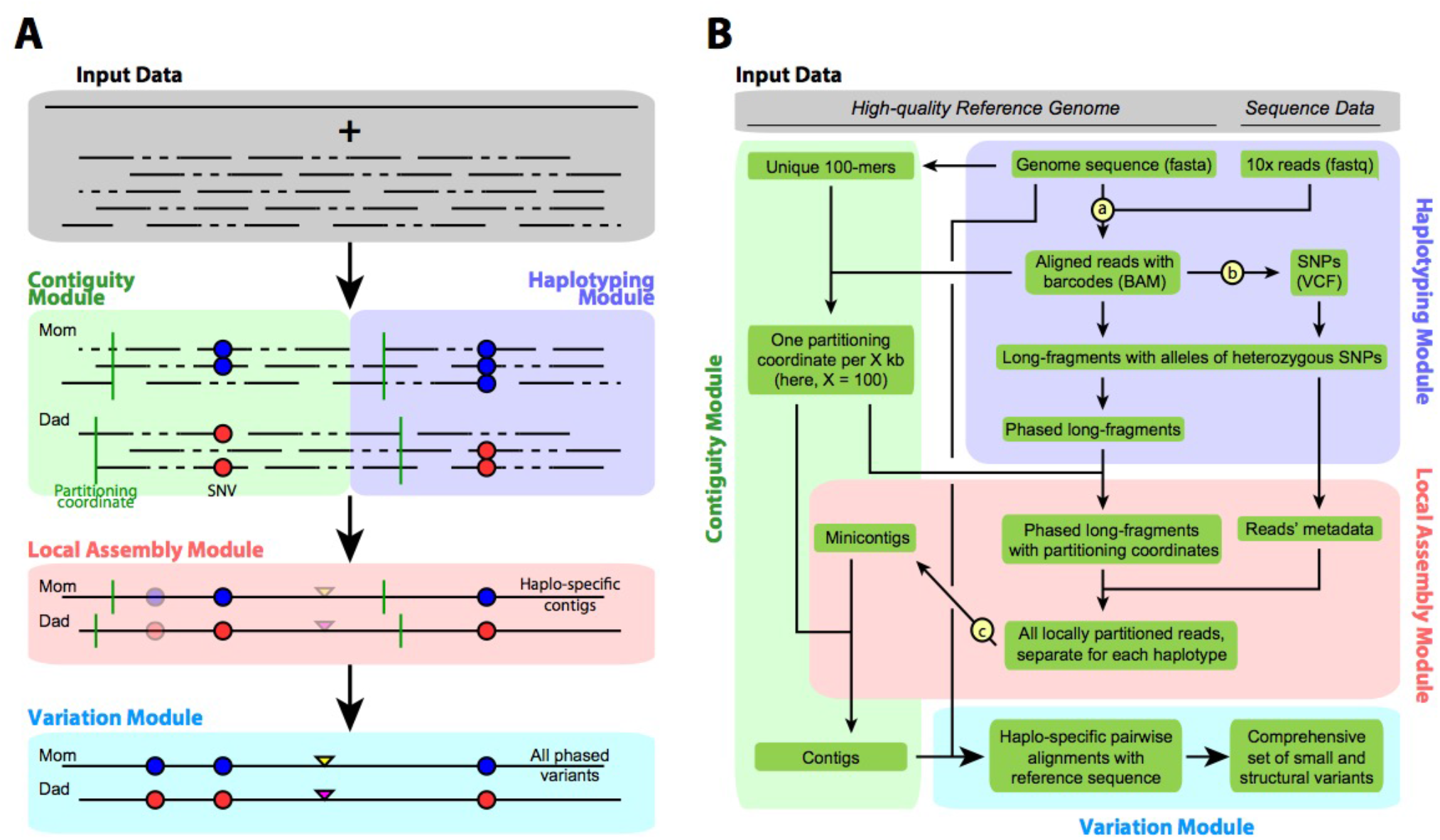
Aquila architecture. Lowercase letters in circles denote existing programs we integrated (a = LongRanger Align, b = FreeBayes, c = SPAdes). (A) Overall architecture. (B) Detailed workflow. Green boxes are data, arrows indicate input and output of a pipeline component. Input data are a high quality reference genome and 10X-based short reads, each with a barcode (not shown). The Haplotyping module produces phased virtual long fragments (by alignment of reads to the reference, SNP detection, and haplotyping) that become part of each read’s record. The Contiguity module produces specific single-base coordinates (‘partitioning points’) in the genome and in the data, where haplotype blocks and reads are cut at a specific single location, and where subsequently assembled minicontigs are rejoined in the end. The Local Assembly module executes assembly of the reads of a specific region, separately for each parental copy. The variation module then discovers, integrates, and infers the phase of all variation.

Each stage has been implemented as a specific module (Figure 1B). In the Haplotyping module, the original long DNA fragments are reconstructed based on barcode-aware alignment of the reads to the reference sequence. In parallel, SNPs are detected based on these same alignments. Fragments are then clustered into either parental bin labeled by each pair of heterozygous SNPs, and a probabilistic model (see methods) is applied to exclude clusters caused by sequencing errors. The clusters are then merged into fewer but larger ones by a greedy recursive algorithm, preserving the separation of parental bins. The resulting clusters contain all reads from a haplotype block of one parent only and are therefore free of allelic variation, greatly simplifying the later assembly steps.

Before assembly is carried out, haplotype blocks whose lengths exceed a user-defined threshold (default = 200kb) are partitioned into smaller chunks (default = 100kb) by the Contiguity module. This step is necessary because a considerable fraction of the data is in large phase blocks (up to the length of an entire chromosome arm) and handing all these reads to the assembler would partially defeat Aquila’s motivation to sidestep the extreme complexity of whole-genome de novo assembly. Thus, the assembly must be broken down into smaller chunks, preferably in places that are highly likely to produce locally correct assemblies, so that contigs from neighboring sub-assemblies can be spliced back together at the partitioning points with high confidence (see Methods; Supplemental Figure S1). The locations of the partitioning points are therefore selected such that (1) they are not within repeats in the reference genome and (2) exhibit the expected read coverage and perfect mapping uniqueness in the abovementioned short-read alignments.

The Assembly module (using SPAdes but any appropriate assembler can be used) produces the complete set of “minicontigs” from the partitions, which are then spliced together to produce final contigs (Supplemental Figure S1). While the emphasis in personal genomes is on producing variant calls (see below), we note that traditional short-read-based methods that go directly from read alignment to variant call do not produce contigs as output. The set of final contigs represents a valuable personal resource in the form of a genome sequence.

The Variation module then aligns the assembled contigs to the reference. All of the small indels and SVs that Aquila reports are detected on the basis of these pairwise alignments. SNPs are also detected but this assembly-based set is merged with the initial set of SNPs identified on the basis of the barcode-aware read alignment performed by the Haplotyping module. The last step in the Variation module is the phasing of all discovered heterozygous variants. The final output is a VCF with all phased variation that also includes all detected variants for which the individual is homozygous compared to the reference sequence, and the small fraction of heterozygous variants that could not be phased.

### Characteristics of Aquila assemblies of six libraries

To explore the performance of Aquila we generated six libraries of 10x linked-read sequencing data from gentle DNA preparations of NA12878 and NA24385 cell lines. The average inferred DNA fragment lengths and their distributions varied among libraries (Supplemental Table S1). We numbered them for each individual according to physical coverage, in ascending order. All of the libraries had approximately 100x Illumina sequencing coverage, except L2 which had 192x. We assembled these libraries with Aquila and compared standard assembly statistics with Supernova 2 assemblies (Zhang et al. 2019b) of the same libraries (Table 1, Supplemental Figure S2). Contiguity, as measured by Contig N50, was generally greater than 100kb; NA50, which is the N50 of contigs after breaking at potential misassemblies by comparison to the reference genome, was in most cases less than 5% lower, indicating few misassemblies. We note that using the reference to identify potential misassemblies is a conservative approach given that it is not the same individual as the sequenced one. Contiguity generally increased as a function of physical coverage (C_F_; L5 producing the best result at C_F_=803x and C_R_=0.08), and was greatest with a weighted fragment length of around 150 kb (Supplemental Figure S2).

**Table 1:** Assembly metrics for the six libraries we built from NA12878 and NA24385 (L1-L6) and a previously published 10x Genomics library (L7). C_F_, physical (‘fragment’) coverage; C_R_, read coverage; C, Raw Coverage >= C_F_ × C_R_. Genome Fraction, percentage of reference genome that is covered by the assembly. Diploid Fraction, percentage of Genome Fraction that is covered by exactly two parental contigs. L5+L6 describes performance for a simple combination of the data from libraries 5 and 6.

| **NA12878** | **Raw**<br>**Coverage**<br>**C (X)** | **C<sub>F</sub>**<br>**(X)** | **C<sub>R</sub>**<br>**(X)** | **Contig**<br>**N50 (bp)** | **Contig**<br>**NA50 (bp)** | **Diploid**<br>**Fraction**<br>**(%)** | **Genome**<br>**Fraction**<br>**(%)** |
| --- | --- | --- | --- | --- | --- | --- | --- |
| **L1_Aquila** | 103 | 123 | 0.41 | 34,759 | 31,645 | 98.1 | 95.45 |
| L1_Supernova |  |  |  | 58,408 | 57,254 | 73.3 | 93.69 |
| **L2_Aquila** | 192 | 334 | 0.27 | 94,699 | 91,662 | 96.7 | 94.43 |
| L2_Supernova |  |  |  | 144,298 | 136,253 | 58.9 | 94.81 |
| **L3_Aquila** | 106 | 958 | 0.07 | 120,963 | 116,438 | 98.7 | 96.17 |
| L3_Supernova |  |  |  | 114,900 | 111,388 | 77.2 | 94.62 |
| **L7_Aquila** | 78 | 243 | 0.32 | 78,309 | 75,378 | 97.1 | 95.32 |
| **NA24385** |  |  |  |  |  |  |  |
| **L4_Aquila** | 100 | 208 | 0.25 | 96,558 | 93,132 | 98.2 | 96.98 |
| L4_Supernova |  |  |  | 99,516 | 96,510 | 79.2 | 95.01 |
| **L5_Aquila** | 100 | 803 | 0.08 | 135,115 | 130,176 | 98.8 | 96.77 |
| L5_Supernova |  |  |  | 129,195 | 125,339 | 78.1 | 94.86 |
| **L6_Aquila** | 100 | 1504 | 0.05 | 125,064 | 120,490 | 98.7 | 96.62 |
| L6_Supernova |  |  |  | 101,236 | 98,364 | 73.4 | 94.69 |
| **L5+L6_Aquila** | 200 | 2307 | 0.07 | 177,974 | 172,358 | 99.1 | 97.06 |

The fraction of the reference genome covered by the assemblies ranged around 95%, indicating that the vast majority of the non-N and not highly repetitive regions of the genome were covered by all libraries assembled by either Aquila or Supernova2. Aquila consistently produced 98% of this fraction in a diploid state, compared to Supernova’s 73-78% (Table 1). This key metric indicates that Aquila produces assemblies that have the potential to support diploid variant detection genome-wide.

Aquila also seamlessly supports combining different libraries, if greater contiguity is desired than is achievable with a single library (Supplemental Figure S3). To illustrate this, we computed the same metrics as for the single libraries for the combination of L5 and L6 from NA24385. Contig N50 increased by 30-40% to 178kb, the fraction of reference genome assembled rose to 97%, and the diploid fraction reached 99% of the assembled genome. While assembly statistics represent an important facet for evaluation of Aquila, the ultimate metric for the usefulness of the approach is how well it detects genetic variation. We therefore evaluated the assemblies for their ability to support detection of SNPs, small indels, and larger structural variants.

### Assembly-based detection of SNPs and small indels

We first evaluated assembly-based SNP and small-indel (<50bp) detection by comparing Aquila’s calls against the Genome in a Bottle (GiaB) benchmark callsets (Zook et al. 2019). The libraries with the best assembly statistics, L3 (from NA12878) and L5 (from NA24385), achieved 97.4% and 97.8% accuracy (F1 metric) for SNPs (Table 2; Supplemental Table S2) and >93% accuracy for the high-confidence set of GiaB small indels (Table 3; Supplemental Table S3). Genotyping errors for the calls that matched GiaB were 0.14%-0.16% for SNPs and 1.61% to 1.90% for high-confidence small indels. The total numbers of assembly-based SNP calls (Supplemental Table S4) were 3,971,444 (L3) and 3,882,869 (L5), compared to the total numbers of FreeBayes-based calls performed on the barcode-aware read alignments of 3,949,721 (L3) and 3,961,684 (L5). Numbers of heterozygotes or homozygotes are also comparable between the two approaches (Supplemental Table S4). We note that Aquila produces numbers of assembly-based SNP calls for these two individuals that are consistent with those previously produced from standard short-fragment Illumina libraries (Wu et al. 2017; Supernat et al. 2018; Li et al. 2018).

**Table 2.** Accuracy of SNP calling, comparing assembly-based calling with two mapping-based approaches on the same libraries’ linked read data, one each from NA12878 (L3) and NA24385 (L5). The benchmark is GiaB v3.3.2. Variant counts and performance scores were generated by RTGtools/hap.py, an Illumina haplotype comparison/benchmarking tool. Longranger calls were executed with “-vcmode=gatk”.

| SNPs |  | True Positives | False Negatives | False Positives | Genotype Mismatch | Precision | Recall | F1 |
| --- | --- | --- | --- | --- | --- | --- | --- | --- |
| L3 (NA 12878) | Aquila | 3,004,501 | 38,282 | 124,074 | 4,730 | 0.960 | 0.987 | 0.974 |
|  | FreeBayes | 3,037,504 | 5,279 | 54,088 | 3,501 | 0.983 | 0.998 | 0.990 |
|  | Longranger | 3,040,701 | 2,082 | 105,854 | 1,621 | 0.966 | 0.999 | 0.983 |
| L5 (NA 24385) | Aquila | 2,989,567 | 39,785 | 93,195 | 4,157 | 0.970 | 0.987 | 0.978 |
|  | FreeBayes | 3,021,814 | 7,544 | 48,477 | 3,899 | 0.984 | 0.998 | 0.991 |
|  | Longranger | 3,026,384 | 2,974 | 104,879 | 1,799 | 0.967 | 0.999 | 0.983 |
| L5+L6 (NA 24385) | Aquila | 2,971,237 | 58,120 | 81,926 | 18,856 | 0.973 | 0.981 | 0.977 |

**Table 3.** Accuracy of small indel calling, comparing assembly-based calling with two mapping-based approaches on the same libraries’ linked read data, one each from NA12878 (L3) and NA24385 (L5).

| Small indels |  | True Positives | False Negatives | False Positives | Genotype Mismatch | Precision | Recall | F1 |
| --- | --- | --- | --- | --- | --- | --- | --- | --- |
| L3<br>(NA 12878) | Aquila | 499,301 | 32,081 | 40,292 | 9,493 | 0.925 | 0.940 | 0.932 |
|  | FreeBayes | 419,344 | 80,354 | 45,977 | 39,636 | 0.903 | 0.839 | 0.870 |
|  | Longranger | 463,732 | 35,966 | 88,431 | 22,513 | 0.843 | 0.928 | 0.883 |
| L5<br>(NA 24385) | Aquila | 476,139 | 26,914 | 35,315 | 7,986 | 0.931 | 0.946 | 0.939 |
|  | FreeBayes | 400,440 | 75,170 | 41,475 | 36,293 | 0.908 | 0.842 | 0.874 |
|  | Longranger | 443,107 | 32,504 | 81,044 | 19,720 | 0.848 | 0.932 | 0.888 |
| L5+L6<br>(NA 24385) | Aquila | 473,895 | 29,158 | 15,724 | 5,335 | 0.968 | 0.942 | 0.955 |

Compared to the GiaB small indel callset, Aquila produces considerably more calls (e.g., 1,007,313 in L3 vs GiaB’s 531,382; Supplemental Table S5). This difference is due to the current incompleteness of the GiaB callset especially among longer small indels, as well as a false-positive rate of the Aquila calls that we cannot rigorously estimate outside of the GiaB regions. The size distribution of Aquila’s small indels matches the size distribution of the GiaB calls very closely, exhibiting the same 2bp periodicity such that insertions or deletions of an even length are more common than those that are one base longer or shorter (Supplemental Figure S4). At lengths above 30bp, where GiaB has very few calls, the Aquila calls continue to exhibit this pattern. The correlations between the Aquila and GiaB distributions of the 1-49 bp indel calls are R^2^=0.997 and 0.998 of the raw counts, and 0.930 and 0.951 for the natural log of counts, for insertions and deletions, respectively.

### Assembly-based detection of structural variants 50bp and greater

Aquila calls ca. 17,000 deletions and ca. 6,000 insertions 50bp and greater in each high quality library (L2, 3, 5, and 6; Supplemental Table S6). The size distributions follow the expected exponential distribution with a peak at ca. 330 bp, which is caused by full-length or nearly full-length Alu elements (Figure 2A). The number of calls is comparable but consistently lower than recent estimates from a comprehensive study that focused exclusively on SVs in a cohort of long-read sequenced individuals, including two haploid samples, in which purpose-driven approaches were applied to achieve high sensitivity of detection of shared SVs (Chaisson et al. 2019; Audano et al. 2019). We do not expect to reach the same level of detection in a single personal genome without the benefit of leveraging several individuals to inform discovery, but we are able to apply several metrics to characterize Aquila’s SV calls (Figure 2B-G).

**Figure 2.**
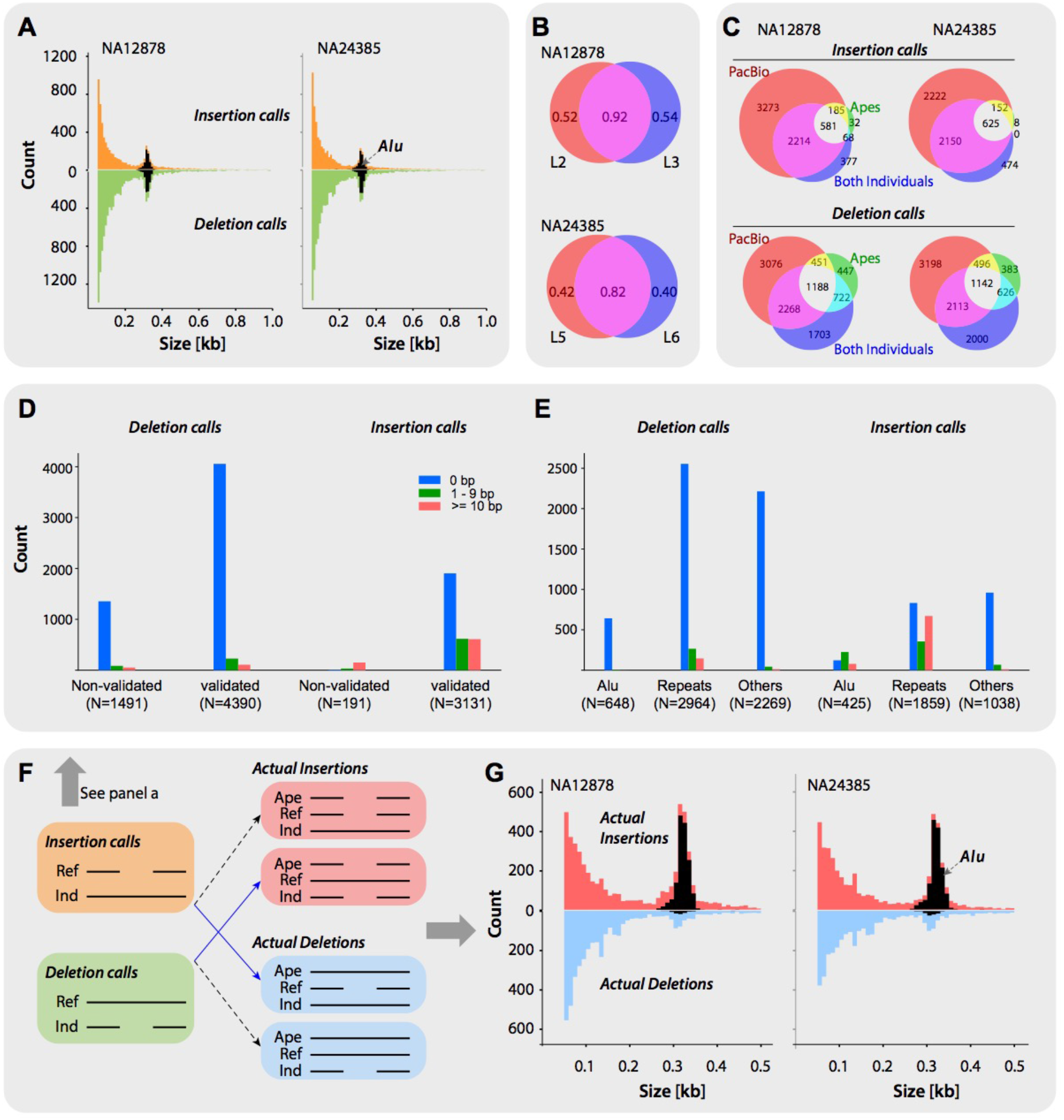
Characteristics and validation statistics of SV calls by Aquila. (A) Size frequency distributions of insertion calls and deletion calls in both individuals (L3 for NA12878 and L5 for NA24385). Black areas represent indels of which at least 80% are a close match to Alu-element consensus sequences. (B) Call validation rates by three validation strategies of two libraries (L2 and L3; L5 and L6) per individual; SVs called in both libraries are in the overlap, flanked by SVs unique to each library. (C) Overlap analysis and comparison of three validation strategies, by call and individual; numbers inside the Venn diagrams are counts of SVs. SVs are validated by: PacBio data from the same individual, (PacBio); the other individual (Both Individuals); in the chimp or orang genome (Apes). Overlaps represent two or more of these criteria fulfilled. (D,E) Comparative precision of SVs present in both individuals, as a function of validation by three validation strategies (D) or sequence class (E). Bar graphs depict counts of SVs that have precisely the same breakpoint coordinates in both individuals (0 bp), that differ by less than 10 (1-9 bp), or that differ by 10 or more (>=10 bp). “Repeats” class includes simple sequence and tandem repeats but not mobile elements; “Other” class includes all SVs that do not overlap more than 80% with Alus and are not part of the Repeats class. (F) Inference of actual molecular mechanism that produced the SV by expanding the alignment between the reference sequence (Ref) and the Individual (Ind) to include chimp or orang sequences; the sequence that matches the ape is the ancestral allele. “Actual insertion” and “Actual deletion” refer to the molecular mechanism that produced the derived allele. Approximately 45% of deletion and 24% of insertion calls are thus ‘inverted’ (blue arrows). (G) Size frequency distributions of actual insertions and actual deletions in both individuals. Black areas represent indels of which at least 80% are a close match to the Alu-element consensus sequence. The peak at around 330 base pairs captures nearly all Alu SVs.

We initially used a combination of three strategies to validate the SV calls in both individuals: First, for both individuals, PacBio data exist that we applied with svviz2 (Spies et al. 2015) to test Aquila’s calls with another data type; second, the two individuals are expected to share a large fraction of SVs, so we performed simple comparisons of the call sets; third, we aligned the SVs and flanking sequence to two high quality ape genomes (Chimp and Orang). Given the complexity of SV calling we expect to have both false negative and false positive calls, which is underscored by the fact that different libraries from the same individual produce strongly overlapping but not fully identical call sets. The calls that are shared between libraries validate (by at least one of the abovementioned three metrics) at 92% for NA12878 and 82% for NA24385 (Figure 2B), whereas calls unique to each library validate at lower rates (40% to 52%). This shows that we do not have perfect sensitivity (because there are many validated calls unique to a single library) and, conversely, that the nonvalidated set of calls unique to a library likely contains false positives.

The number of SVs validated by svviz2/PacBio is higher than those validated by the other two approaches (Figure 2C), since the PacBio data is from the same individual as the respective assembly. However, there are many SVs that are not validated by PacBio but that are also called in the other individual or are validated by comparison to ape sequences. This effect is greater for deletion calls than for insertion calls (Figure 2C), which suggests that insertions may be called at higher specificity and lower sensitivity than deletions. This interpretation is consistent with a predicted shortcoming of the current implementation of Aquila, which because of its reliance on the reference sequence to identify reads for assembly has decreasing power to assemble insertions as their size increases.

We also assessed the consistency of the breakpoints of the calls that are shared between the two individuals (Figure 2D,E). We binned the SVs based on the size differences between the calls in NA12878 and NA24385, as a function of validation by the other two approaches (Figure 2D) or as a function of sequence type (Figure 2E). Overall, the vast majority of calls have precisely the same breakpoints in both individuals. Deletions or validated calls have better precision than insertions or nonvalidated calls, and SVs in repeats have worse precision than nonrepetitive sequences.

### Genome in a Bottle Benchmark Comparison

During the course of this work, the first GiaB SV benchmark, v0.6, was released. It is based on the HG19 reference sequence and is specific to NA24385, with 9397 SVs greater than 50 bp in the call set. Accuracy (F1 metric) of Aquila calls, assuming all GiaB calls are correct, ranges from 52% to as high as 87% depending on the type of variant and the location in which it occurs (Table 4). This compares favorably with results obtained from Supernova assemblies of the same libraries (Zhang et al., 2019b). In general, Aquila’s recall performance is better than precision, which is likely due to a combination of a rate of actual false positives called by Aquila and an unknown number of false negatives in the GiaB callset.

**Table 4.** Overall benchmarks of SVs from L5, L6 and L5+L6 by Aquila. SVs were called from the NA24385 assemblies and compared to GIAB NIST_SVs_Tier1_v0.6 (bed file HG002_SVs_Tier1_v0.6_chr.bed). SV counts and performance scores were generated by Truvari (parameters -p 0.1 -P 0.1 -r 200 --passonly). no TR > 100, without SVs if at least 20% of the reference bases are tandem repeats at least 100bp long; no TR any size, without SVs if at least 20% of the reference bases are in tandem repeats of any size. Numbers in parentheses (not-in/in break) are counts of SVs outside or within assembly breaks.

| SVs |  | True<br>Positives<br>(not-in/in<br>break) | False<br>Negatives<br>(not-in/in<br>break) | False<br>Positives<br>(not-in/in<br>break) | Precision<br>(not in<br>break) | Recall<br>(not in break) | F1<br>(not in break) |
| --- | --- | --- | --- | --- | --- | --- | --- |
| L5 | Total | 6,197<br>(5,631/566) | 3,200<br>(1,424/1,776) | 7,776<br>(6,181/1,595) | 0.443<br>(0.477) | 0.659<br>(0.798) | 0.530<br>(0.597) |
|  | no TR ><br>100bp | 3,900<br>(3,578/322) | 1,636<br>(445/1,191) | 3,806<br>(3,046/760) | 0.506<br>(0.540) | 0.704<br>(0.889) | 0.589<br>(0.672) |
|  | no TR any<br>size | 1,546<br>(1,429/117) | 916<br>(297/619) | 466<br>(398/68) | 0.768<br>(0.782) | 0.628<br>(0.828) | 0.691<br>(0.804) |
| L6 | Total | 6,086<br>(5,551/535) | 3,311<br>(1,460/1,851) | 8,021<br>(6,264/1,757) | 0.431<br>(0.470) | 0.648<br>(0.792) | 0.518<br>(0.590) |
|  | no TR ><br>100bp | 3,830<br>(3,515/315) | 1,706<br>(471/1,235) | 3,889<br>(3,036/853) | 0.496<br>(0.537) | 0.691<br>(0.882) | 0.577<br>(0.667) |
|  | no TR any<br>size | 1,527<br>(1,428/99) | 935<br>(301/634) | 430<br>(365/65) | 0.780<br>(0.796) | 0.620<br>(0.826) | 0.691<br>(0.811) |
| L5+L6 | Total | 6,434<br>(5,983/451) | 2,963<br>(1,416/1,547) | 5,577<br>(4,404/1,173) | 0.536<br>(0.576) | 0.685<br>(0.809) | 0.601<br>(0.673) |
|  | no TR ><br>100bp | 4,065<br>(3,803/262) | 1,471<br>(456/1,015) | 2,848<br>(2,256/592) | 0.588<br>(0.628) | 0.717<br>(0.893) | 0.646<br>(0.737) |
|  | no TR any<br>size | 1,671<br>(1,574/97) | 791<br>(294/497) | 223<br>(185/38) | 0.882<br>(0.895) | 0.662<br>(0.842) | 0.756<br>(0.867) |

### Inference of derived alleles

SV calls are labeled with respect to the reference sequence as ‘insertion’ or ‘deletion’, but the molecular mechanism that generated the SV may be the opposite of the call because the reference sequence is a random sample of ancestral and derived alleles from the population. Where the reference carries a derived allele, the actual molecular mechanism that generated the SV is the opposite of the call because the assembly carries the ancestral allele. For those SVs with alignments to ape genomes, the SV allele matching the ape sequence is highly likely to represent the ancestral state, and the other allele is derived, allowing inference of the actual molecular mechanism that generated it (Figure 2F). Classifying the SVs accordingly (“actual” insertion or deletion) causes a striking shift in the size distributions of insertions and deletions in both individuals (Figure 2A,G), in which the vast majority of SVs that overlap Alu repeat sequences are now revealed to be actual insertions (Figure 2G). This provides empirical evidence that the classification into ancestral and derived alleles is largely correct, as Alus are known to insert as full length sequences, whereas partial Alus are degenerate copies that arise later in evolution from deletions whose breakpoints can be anywhere in the element.

### Genome-wide distribution and phasing of all variation

To interrelate the different types of variation detected by Aquila and to ask whether there were any obvious biases we divided the genome into bins of 250kb and quantified tandem repeat content, contig density, and variation content by genotype from the assembly of L3 (Figure 3). Contig density, which over the vast majority of the genome is exactly 2 because of the diploid nature of the assemblies, does not correlate with any variation and only weakly with repeat content. Repeat content correlates weakly with the number of SVs. As expected, numbers of SNPs and very small indels correlate strongly when the genotype is the same (heterozygous or homozygous), and weakly with larger small indels, which in turn correlate weakly with SVs. Overall, the correlation patterns do not reveal any large-scale biases in variation discovery. We also note that the fraction of variants that are heterozygous varies over a narrow range across all types and sizes of detected variation (Supplemental Figure S5), again revealing no obvious biases.

**Figure 3.**
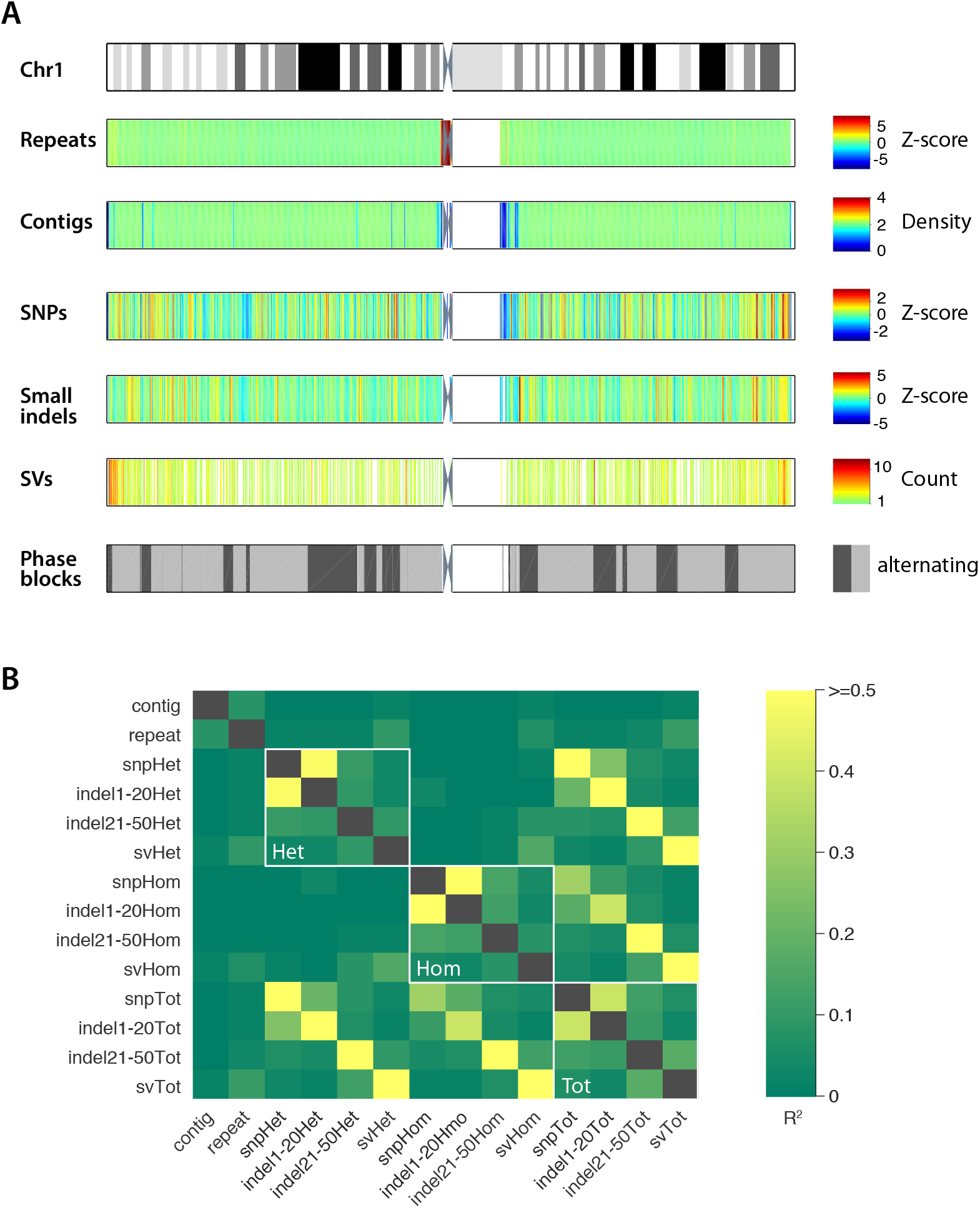
Local distribution of all types of variation detected, aggregated across 250kb intervals. (A) Top, ideogram of Chromosome 1 showing metaphase banding pattern. Tracks below are assembly and variation features where white represents no data. Repeats, Z-score of TRF-detected repeat density. Contigs, average number of contigs per base pair. SNPs, Z-score of SNP number. Small indels, Z-score of the number of indels 1-50bp. SVs, number of SVs >=50bp in each 250kb bin. Phase blocks, each phase block is a grey rectangle, with alternating light and dark indicating neighboring phase blocks. (B) Genome-wide correlation (R^2^) among all pairs of variation types by genotype, contig density, and repeat density.

The last step in Aquila’s pipeline is a final integrative phasing of all of the discovered heterozygous variation on the basis of the phase blocks obtained with heterozygous SNPs in the Haplotyping module (Table 5). Depending on the library, between ca 1.7 and 2 million SNPs were initially phased (Supplemental Table S7). Because the parental genotypes are known for the two individuals we could quantify the phasing error, which is dominated by switch errors that involve a single, presumably incorrectly genotyped, SNP (Table 5). Long switch errors are quite rare, comparing favorably with previous work using 10x data and a variety of phasing algorithms (Shajii et al. 2018; Marks et al. 2019). Because SNPs are the densest type of polymorphism in human genomes, phasing other variants on the basis of these is feasible. Instead of probabilistic imputation, however, Aquila performs straightforward inference by matching the assembly-based SNP calls with those of the Haplotyping module and then simply inferring the correct phase (Supplemental Figure S6). In total, Aquila added ca. 1 million heterozygous variants, including 0.5 – 0.8 million previously unphased SNPs, ca. 0.5 million small indels, and ca. 10,000 heterozygous SVs in the best four libraries (Supplemental Table S7).

**Table 5.**
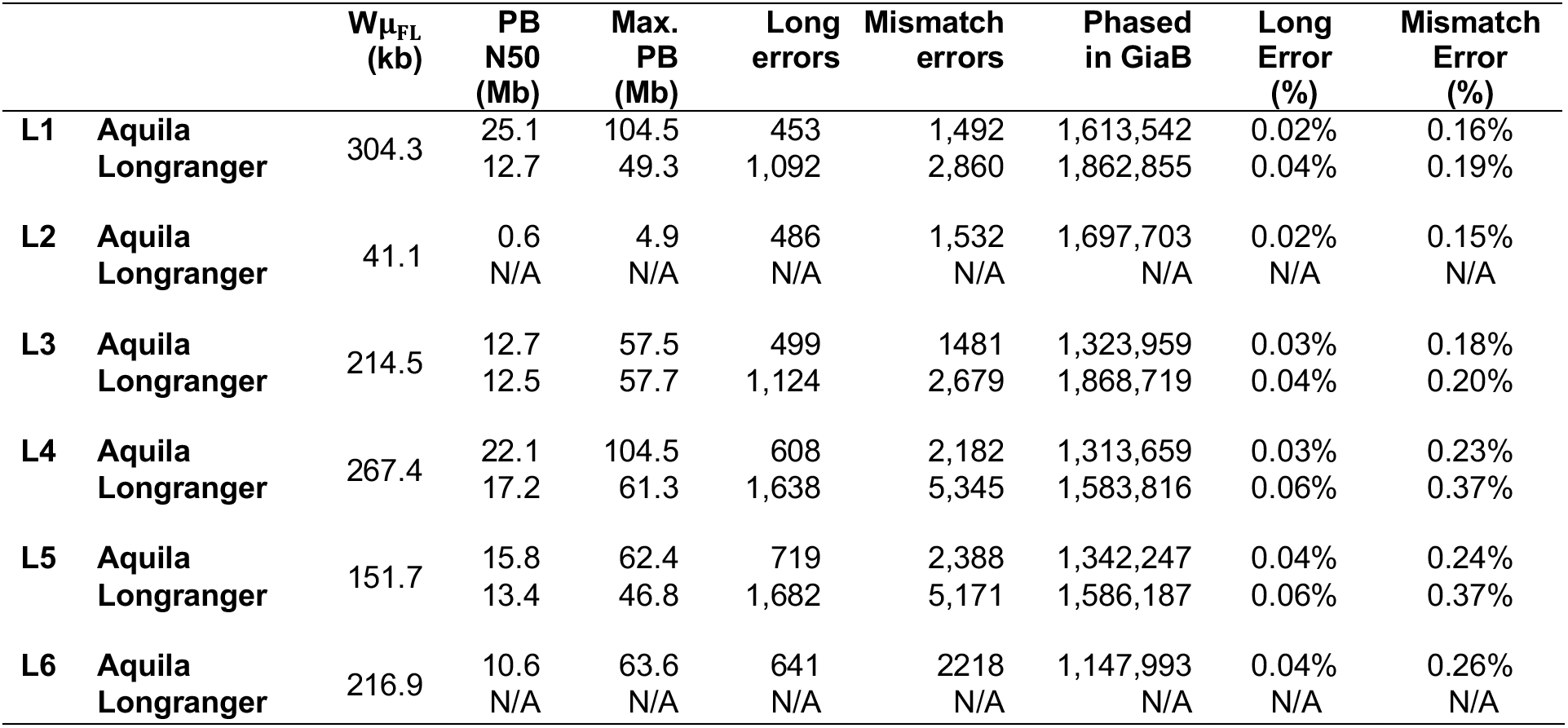
Phasing information and accuracy for each library from Aquila versus Longranger. W_*μFL*_, weighted fragment length of the library; PB N50, phase blocks length N50; Max. PB, maximum phase block length; Phased in GiaB, number of phased SNPs overlapping with callset v3.3.2. N/A, not applicable because Longranger could not complete runs in wgs mode.

| | | $W\mu_{FL}$<br>(kb) | PB<br>N50<br>(Mb) | Max.<br>PB<br>(Mb) | Long<br>errors | Mismatch<br>errors | Phased<br>in GiaB | Long<br>Error<br>(%) | Mismatch<br>Error<br>(%) |
| --- | --- | --- | --- | --- | --- | --- | --- | --- | --- |
| L1 | Aquila<br>Longranger | 304.3 | 25.1<br>12.7 | 104.5<br>49.3 | 453<br>1,092 | 1,492<br>2,860 | 1,613,542<br>1,862,855 | 0.02%<br>0.04% | 0.16%<br>0.19% |
| L2 | Aquila<br>Longranger | 41.1 | 0.6<br>N/A | 4.9<br>N/A | 486<br>N/A | 1,532<br>N/A | 1,697,703<br>N/A | 0.02%<br>N/A | 0.15%<br>N/A |
| L3 | Aquila<br>Longranger | 214.5 | 12.7<br>12.5 | 57.5<br>57.7 | 499<br>1,124 | 1481<br>2,679 | 1,323,959<br>1,868,719 | 0.03%<br>0.04% | 0.18%<br>0.20% |
| L4 | Aquila<br>Longranger | 267.4 | 22.1<br>17.2 | 104.5<br>61.3 | 608<br>1,638 | 2,182<br>5,345 | 1,313,659<br>1,583,816 | 0.03%<br>0.06% | 0.23%<br>0.37% |
| L5 | Aquila<br>Longranger | 151.7 | 15.8<br>13.4 | 62.4<br>46.8 | 719<br>1,682 | 2,388<br>5,171 | 1,342,247<br>1,586,187 | 0.04%<br>0.06% | 0.24%<br>0.37% |
| L6 | Aquila<br>Longranger | 216.9 | 10.6<br>N/A | 63.6<br>N/A | 641<br>N/A | 2218<br>N/A | 1,147,993<br>N/A | 0.04%<br>N/A | 0.26%<br>N/A |

## Discussion

We introduce a new method, Aquila, which uses a high-quality reference sequence to perform assembly in small chunks for both haplotypes, producing a diploid personal genome sequence from a single data type, Illumina/10X linked reads. We test Aquila’s performance on several libraries from two standard individuals and show that it produces high quality diploid assemblies. The assemblies enable comprehensive discovery of SNPs, small indels, and SVs, on the basis of pairwise alignments to the reference genome. We show that Aquila produces overall better results than de-novo assembly from Supernova (Zhang et al. 2019b), for all types of variants. Accuracy of variant discovery, as evaluated against GiaB benchmark sets, retains some characteristics of standard Illumina short-fragment sequencing, with higher accuracy for smaller variants than for longer ones. Long-range phasing is highly accurate with low switching error.

Aquila is the first approach that effectively leverages the strengths of 10X/Illumina sequencing to enable comprehensive variation discovery and phasing in personal genomes at a reasonable cost. Compared to standard Illumina sequencing, the increased cost of the 10X sample prep and the generation of deeper sequence data is justified by (1) the much greater power to detect insertions and deletions, and (2) the genome-wide phasing of all heterozygous variants. Compared to ONT and standard PacBio, the superior base-pair level accuracy of Illumina sequencing ensures more accurate SNP and small indel detection as well as SV breakpoint determination; and while PacBio CCS appears to have the potential to enable highly accurate genome sequencing (Wenger et al. 2019), it remains to be seen whether it performs substantially better than 10X/Illumina/Aquila to justify the additional cost, particularly for cohort studies. Compared to ensemble approaches that computationally integrate multiple data types with complementary strengths (for example, combining Illumina, long-read, and BioNano data), 10X/Illumina/Aquila is far less complex to manage in the laboratory and offers a simpler computational approach.

In the ecosystem of solutions for human whole genome sequencing, 10X/Illumina/Aquila therefore fills what we believe is the most important niche: a diploid and phased personal genome for accurate and comprehensive discovery of SNPs, small indels, and SVs in all but the most complicated regions of the human genome. It represents the first generation of approaches that drive toward laboratory and computational efficiency and simplicity by using a single data type and leveraging the considerable amount of information present in the human reference sequence. Until de novo assembly on the basis of highly accurate very-long-read data is shown to be cost-effective, reference-assisted approaches that partition the genome into smaller assembly problems are likely to prevail.

Further improvements of the approach we take here fall into two categories: those for which the nature of current linked reads data is inherently limited and will require technological advances and those in which future implementations of Aquila will produce better results. For example, it is unlikely that linked reads data will support assembly and resolution of recent segmental duplications or long repetitive sequences. However, the current dropoff in sensitivity to detect insertions beyond 500 bases will be addressed by improving the inclusion of ambiguously mapping reads into the assembly process. Similarly, improvements to assembly contiguity will increase accuracy of variants in repetitive sequences, which are currently enriched in assembly breaks. Future improvements will also center on better detection of long insertions and contig breakpoint assembly. Although developed for 10X/Illumina data, Aquila’s architecture may be used in the future for computational approaches that use data from other sources. In its current form, it is already applicable to any study that requires better indel detection than what is achievable with standard Illumina sequencing.

## Methods

Aquila is organized in four conceptual modules that correspond to the following python steps (Figure 1): Aquila_step1.py: Haplotyping module + contiguity module (first part: partitioning). Aquila_step2.py: Local assembly module + contiguity module (second part: concatenation). Variation Module: Aquila_assembly_based_variants_call.py and Aquila_phasing_all_variants.py.

### Pruning unreliable variants (Haplotyping module)

Accurate haplotyping requires filtering out incorrectly genotyped variants and false positives due to sequencing error. We performed an empirical analysis for 10x data to investigate the alternate allele frequency (*R*_*alt*/*ref*_) and coverage per variant (*d*_*var*_) that could be used as metrics to find erroneous calls. Allele frequency (*R*_*alt*/*ref*_ >= 0.25) was used for a cutoff, and a 2-tailed percentile cutoff was used for coverage per variant (*10%*avg_cov* <=*d*_*var*_ <= *90%*avg_cov, avg_cov*: average read coverage per variant). The haplotyping algorithm was further improved by sacrificing a small amount of low-confidence heterozygous variants. SNP quality (*13* by default) is the final free parameter used to prune variants.

### Inference and phasing of original long-fragments (Haplotyping module)

All DNA fragments are first reconstructed by aligning short reads to the human genome reference (Hg38) by a barcode-aware alignment strategy (‘*Longranger align’,* https://support.10xgenomics.com/genome-exome/software/pipelines/latest/installation). Aquila then sorts all reads by barcodes and positions, and collects the reads with the same barcode to reconstruct each fragment. There is a threshold to differentiate two molecules with the same barcode when the distance between two successive reads with the same barcode is larger than 50kb (50kb by default, free parameter). After reconstructing all fragments, Aquila assigns the alleles of heterozygous SNPs to each fragment by scanning the reads belonging to each fragment and comparing to a VCF file generated by FreeBayes. At a heterozygous locus “*0*” is the reference allele and “*1*” is the alternate allele.

For each pair of heterozygous variants, if the even parity was correct where one haplotype supported “*00*”, and the other haplotype supported “*11*”, the odd parity must then have been caused by a sequence error with some fragments supporting “*01*”, and other fragments supporting “*10*”, and vice versa. In rare scenarios, the fragment could have two sequencing errors when the even/odd parity was correct, but the fragment supported the complementary haplotype (e.g., the haplotype is ““*00*””, and the fragment supports “*11*”). For each fragment with at least two heterozygous SNPs, Aquila records all neighboring pairs of heterozygous variants. It then applies a Bayesian model (see below) to evaluate if even or odd parity is correct, and the clusters with the parity caused by sequencing error are excluded from further steps. Importantly, the excluded clusters are due to the variants caused by sequencing errors, not the molecules themselves, which means if these molecules still contain other pairs of heterozygous variants with consistent haplotype with the correct parity, they are still used for haplotyping.

Aquila then performs a recursive clustering algorithm in two haplotypes to aggregate bigger clusters/phase blocks. Two clusters are merged if the number of molecules in both of them supporting the same haplotype exceeds a threshold. This threshold is set to 3 by default, which corresponds to a merging error percentage ≤ *((1−p*_*1*_)*(1−p*_*2*_))^*3*^, for each pair of variants, if each variant matched the true variant with probability *p*_*1*_ and *p*_*2*_, respectively. Aquila sorts all pairs of clusters by the positions of reads of all molecules in each cluster. When two locally successive clusters are merged into one single cluster, the corresponding clusters of the other haplotype are merged too. The resulting pairs of clusters are sorted again for the next iteration. The sorting algorithm complexity is ~ *O(N*_*var*_*logN*_*var*_), where *N*_*var*_ is the total number of heterozygous variants. Aquila performs clustering recursively until no more clusters can be merged based on the supporting threshold.

The result of this step are pairs of clusters where each pair corresponds to one diploid phase block. For each phase block, Aquila then performs haplotype construction and extension when each heterozygous variant is supported by all the molecules that cover it. When there are multiple molecules supporting inconsistent genotypes for a variant, that variant is excluded from further steps. To further extend the phase blocks, Aquila similarly performs recursive clustering when two phase blocks have a number of overlapping variants greater than a certain threshold. The threshold is set to 5 by default so that the merging error due to sequencing error p is ≤ *p*^*5*^. When no more phase blocks can be merged the process has converged.

In the final step, all the molecules/fragments with at least two heterozygous variants are assigned to a phase block based on the variants in the final phase block, and a maximum likelihood estimation is applied. Given a haplotype *H* and a molecule *M*, we apply the theta function *θ*(*H*_*i*_, *M*_*i*_) = 1 if *H*_*i*_ = *M*_*i*_ and *0* otherwise. Given *p*_*i*_, the probability the allele call at variant *i* in molecule *M* is correct, the likelihood of observing molecule *M* is:

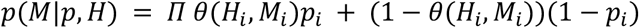

Similarly, given the complementary haplotype *H*_*c*_ and molecule *M*, a theta function gets the value *θ*(*H*_*ci*_, *M*_*i*_) = 1 if *H*_*ci*_ = *M*_*i*_ and *0* otherwise. The likelihood of observing molecule *M* is: *p*(*M*|*p*, *H*_*c*_) = *Π θ*(*H*_*ci*_, *M*_*i*_)(1 − *p*_*i*_) + (1 − *θ*(*H*_*ci*_, *M*_*i*_))*p*_*i*_. To assign the final phase block to each molecule *M*, Aquila needs to find the haplotype *j* in the final phase blocks that meets 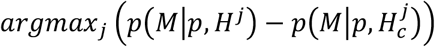.

### Probability model to determine correct joint key (Haplotyping module)

At each position, the genome has two complementary “keys” matching the true two haplotypes: *00*, *11* (even parity) or *01*, *10* (odd parity). Each variant matches the true variant with probability *p*_*1*_ and *p*_*2*_, respectively. The probability that a sequence key will have the correct parity is *p*_*c*_ = *p*_*1*_*p*_*2*_+(1−*p*_*1*_)(1−*p*_*2*_), since both variants could match the true variants or both variants are called wrong. Let *N* be the number of sequences observed that have both variants, and *k* be number of sequences with key of even parity, and *(N − k)* be the number of sequences with key of odd parity. Using Bayes’ theorem to test *P(B | A)*:

Where *A*: *k* out of *N* molecules have keys with even parity, *B*: true key is even parity.

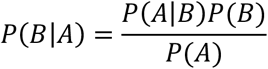

Aquila will accept even parity is correct if *P*(*B* | *A*) exceeds a significance level (e.g. > *0.99*), where:

*P*(*B* | *A*) is the probability of the true key being even parity given *k* out of *N* molecules have keys with even parity.
*P*(*A* | *B*) is the probability of *k* out of *N* molecules having keys with even parity given the true key is even parity, which is 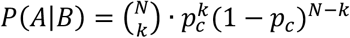.
*P(A)* is the probability of *k* out of *N* molecules have keys with even parity, which is 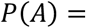 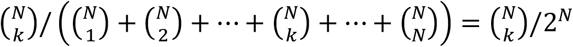.
*P(B)* is the probability of even parity being correct, *P (B)* = *1*.

### High-confidence partitioning point profile generation (Assembly module)

Large phase blocks (free parameter: *block_threshold = 200kb by default*) are cut into multiple small chunks of a specific length (free parameter: *block_len_use = 100kb by default*) based on a high-confidence partitioning point profile. This is done to make assembly faster and more tractable, by avoiding too many reads being given to the assembler. This profile is generated based on three criteria: 1. Expected reads coverage (*C*), 2. Expected physical coverage (*C*_*F*_), 3. *100*-mer uniqueness. Read depth and fragment/physical coverage for each position is calculated after reconstructing all the fragments. The 100-mer uniqueness file (Karimzadeh et al. 2018) for hg38 processed and included in Aquila, was downloaded from http://hgdownload.soe.ucsc.edu/gbdb/hg38/hoffmanMappability/k100.Unique.Mappability.bb Each locus/position is defined to be a high-confidence partitioning point if *C*_(partitioning point) >_ *C*_*(average)*_ *0.8, *C*_*F (partitioning point)* >_ *C*_*F (average)*_ *0.8, and *locus ∈ 100-mer* uniqueness. Aquila uses these high-confidence partitioning points in the profile as reference points to partition reads before assembly (see next section) and later to reconnect the resulting mini-contigs into contigs (Supplemental Figure S1).

### Local assembly within small diploid chunks and stitching contigs (Assembly and Contiguity modules)

For each phase block (Supplemental Figure S1), Aquila records its original long molecules and their corresponding short reads. Large phase blocks are cut into small chunks (see previous step) to perform local assembly with SPAdes (Bankevich et al. 2012) within each phase block for both haplotypes separately. (SPAdes is included in the Aquila package.) Those resulting minicontigs from neighboring small chunks that are bounded by the same partitioning point are concatenated. 99% of partitioning points met this criterion. For each concatenating iteration, the previous concatenated contig is used for the next iteration of concatenation. At the end of this step, Aquila has generated contigs for both haplotypes in each original phase block. The algorithm complexity is ~ *N*_*chunks*_*O*(*T*_*onechunk*_), where *N*_*chunks*_ = number of small chunks and *T*_*onechunk*_ = time for finishing assembly of one small chunk.

### Variation detection for assembled contigs (Variation Module)

To generate single nucleotide polymorphisms (SNPs), small Indel, and structural variant (SV) calls from the *de novo* assemblies, Aquila uses the contig file of the haplotype 1 and haplotype 2 of each phase block. Minimap2 (Li 2018) and paftools (https://github.com/lh3/minimap2/tree/master/misc) are integrated and applied to call variants from each (haploid) contig (“-cx asm5 ‒cs” is applied for minimap2, and “-l 1 -L 1 -q 20” is applied to paftools). For contig alignments and variant selection, mapping quality (>=*20*) is chosen to produce variant candidates. Finally, to generate SNPs, if both haplotypes cover the alternate allele, it is defined as homozygous; if one haplotype covers the reference allele and the other haplotype covers the alternate allele, it is defined as heterozygous.

To generate small Indels and SVs, variant candidates from each haplotype are compared against each other to infer zygosity. To achieve that, heterozygous variants are defined if one haploid assembly contains alternate allele(s) and the other haploid assembly contains reference allele(s). Homozygous variants are defined if both haploid assemblies contain alternate allele(s). For compound indel/SV, we split them into two heterozygous variants. Check “--all_regions_flag = 1” for “Aquila_assembly_based_variants_call.py” in GitHub to perform these analyses.

### Phasing inference (Variation Module)

The initial phased SNPs from the Haplotyping module provide the scaffold on which all other heterozygous variants that are discovered by the Variation module are phased (Supplemental Figure S6). For example, consider the case of one assembled SNP in a phase block, “G|A” or “1|0” (where “A” is the reference allele, and “G” is the alternate allele), and the other neighboring assembled SNP in the same phase block, “C|T” or “0|1” (where “C” is the reference allele, and “T” is the alternate allele): these two phased SNPs have the same genotype and phase in a phase block from the haplotyping module. Therefore, Aquila places the SV into the haplotype that is in the same phase. This is done for all heterozygous variants discovered by the Variation module.

## Supporting information

SupplementalFile_Aquila

## Software availability

Aquila can be found at https://github.com/maiziex/Aquila. For easy installation, install through Bioconda by “conda install aquila”. Version 1.0.0 was used to generate the results in this paper.

## Data availability

The raw sequencing data can be downloaded in the Sequence Read Archive and its BioProject accession number is PRJNA527321(Zhang et al. 2019a). Assemblies and VCFs can be found at http://mendel.stanford.edu/supplementarydata/zhou_aquila_2019/. We will be submitting raw sequence data and assemblies to NCBI’s SRA and Assembly databases.

## Acknowledgements

This research was supported by the Joint Initiative for Metrology in Biology (JIMB; National Institute of Standards and Technology). We would like to thank Noah Spies, Justin Zook, and Marc Salit for informative discussions.

