## SupplementalFile_Aquila for "Aquila: diploid personal genome assembly and comprehensive variant detection based on linked reads"

### Supplemental Methods and Results

#### Contig quality assessment

We used QUAST (Gurevich et al., 2013) to generate various assembly metrics such as N50 and NA50. *--extensive-mis-size* 1000 was applied as the lower threshold of the relocation size.

#### Aquila computation cost

Aquila's four modules vary in their memory requirements and are flexible with respect to computing architecture, allowing jobs to be run in parallel on a cluster or serially in a large-memory machine. Generally the run time is considerably longer on the large memory machine. On a cluster, jobs are parallelized by chromosome or a combination of chromosomes that minimizes the number of nodes required. Modules 1 to 3 use 23, and module 4 uses 10 hours of wall clock time respectively on a current-generation standard compute cluster (23 nodes with 128Gb RAM each, 2 CPUs each, 10-cores per CPU, i.e. 240 cores). Check tables of computation cost in GitHub for more details.

#### Aquila output

Aquila outputs an overall contig file “Aquila_contig.fasta”, and separately for each chromosome “Aquila_Contig_chr*.fasta”. After performing assembly-based variant calling in the Variation Module, one contig file for each haplotype is generated: “Aquila_Contig_chr*_hp1.fasta” and “Aquila_Contig_chr*_hp2.fasta”. For each contig, the header (for example “*>36_PS39049620:39149620_hp1*”) follows the format contig number (“*36*”), phase block start coordinate (“*PS39049620*”), phase block end coordinate (“:*39149620*”), and haplotype number (“*hp1*”). Within the same phase block, the haplotype number “*hp1*” and “*hp2*” are arbitrary for maternal and paternal haplotypes. For some contigs from large phase blocks, the headers are much longer and complex, for an instance, *“>56432_PS176969599:181582362_hp1_ merge177969599:178064599_hp1-177869599:177969599_hp1*”. “*56*” denotes contig number, “*176969599*” denotes the start coordinate of the final big phase block, “*181582362*” denotes the end coordinate of the final big phase block, and “hp1” denotes the haplotype “*1*”. “*177969599:178064599_hp1*” and “*177869599:177969599_hp1*” mean that this contig is concatenated from minicontigs in small chunk (start coordinate: *177969599*, end coordinate: *178064599*, and haplotype: *1*) and small chunk (start coordinate: *177869599*, end coordinate: *177969599*, and haplotype: *1*).

#### Combining multiple libraries of linked-reads into a single assembly

Each library of 10x linked-read sequencing data uses the same set of barcodes, which makes the combination of multiple libraries at the stage of raw fastq reads difficult. To combine multiple libraries, Aquila reconstructs the original fragments with their corresponding reads of each library separately, after which barcodes are not required any more. All the DNA fragments from multiple libraries, and their corresponding reads, are then combined to perform haplotyping and all subsequent steps. The molecule haplotyping algorithm is efficient and makes haplotyping fragments for multiple libraries fast. The algorithm complexity is linear with the number of libraries, ~ $N_{libs}O(T_{onelib})$, where $N_{libs}$ = number of libraries and $T_{onelib}$ = time for finishing assembly of one library.

#### Validation and evaluation methods for variations

To validate the SNP calls from the assembled contigs, we used the GiaB benchmark call sets for both NA12878 and NA24385. They included 3,084,732 SNPs (1,0210,585 homozygous and 1,874,147 heterozygous) for NA12878, and 3,076,552 SNPs (1,180,678 homozygous and 1,895,874 heterozygous) for NA24385. The benchmark allowed us to calculate both the sensitivity and the genotype accuracy of each SNP called from assemblies. We performed the same analysis on the small indel calls with the small indel callset 0.6 from GiaB (NA12878: ftp://ftp-trace.ncbi.nlm.nih.gov/giab/ftp/release/NA12878_HG001/latest/GRCh38/, NA24385: ftp://ftp trace.ncbi.nlm.nih.gov/giab/ftp/release/AshkenazimTrio/HG002_NA24385_son/latest/ GRCh38/). For SV validation we applied svviz2 (https://github.com/nspies/svviz2) with PacBio reads (NA12878: ftp://ftp-trace.ncbi.nlm.nih.gov/giab/ftp/data/NA12878/NA12878_PacBio_MtSinai/ and NA24385: ftp://ftp-trace.ncbi.nlm.nih.gov/giab/ftp/data/AshkenazimTrio/HG002_NA24385_son/PacBio_MtSinai_NIST/).

We classified SVs into three categories: Alu, Tandem Repeats, and Other. We selected SVs from 250-350bp for classification as Alu elements by alignment to the AluY consensus sequence, and only SVs that matched at 80% or more were labelled as Alus. We used Tandem Repeats Finder (trf, (Benson, 1999)) to label repeats/non-repeats for all non-Alu-SVs, and the repeat percentage was calculated. Sequences that did not meet the Alu alignment criterion and did not contain a tandem repeat were labeled "Other".

For assemblies of different libraries, the different assemblies could identify different SV candidates. To analyze the overlapping and unique SV calls of three libraries of the same individual, we also merged each SV from different libraries. We applied the same merging criteria when we called homozygous/heterozygous SV from two haplotypes. If the coordinates of two deletions overlapped we defined them to be the same deletion, and if the breakpoints of two insertions were within *20bp*, we defined them to be same insertion.

#### Ancestral analysis

For each called SV, we extracted the left and right flanking 500bp, and aligned them to the Orangutan and Chimpanzee references. For deletion calls, if the end coordinate of the left flanking sequence was within 2bp of the start coordinate of the right flanking sequence, this SV was defined as “Ins_Ref” (insertion on reference - an actual insertion). If the distance between the end coordinate of the left flanking sequence and the start coordinate of the right flanking sequence was the approximate length of the called deletion (*0.9*SV_size - 1.1* SV_size*), this SV was defined as “Del_Tar” (deletion on target - an actual deletion). For insertion calls, if the end coordinate of the left flanking sequence was within 2bp of the start coordinate of the right flanking sequence, this SV was defined as “Ins_Tar” (insertion on target - an actual insertion). If the distance between the end coordinate of the left flanking sequence and the start coordinate of the right flanking sequence was the approximate length of the called deletion (0.9*SV_size - 1.1* SV_size), this SV was defined as “Del_Ref” (deletion on reference - an actual deletion).

We performed multiple sequence alignments (for each SV and its aforementioned flanking regions annotated as “Ins_Ref”, “Del_Tar”, “Ins_Tar” and “Del_Ref”) with Muscle (Edgar, 2004).

#### Comparison of Genotype Frequencies

We asked whether segregation of the SV alleles behaves as expected. For each locus, we have two chromosomes each from the sequenced individuals and one from the reference genome, for a total of 5, and therefore 18 possible combinations of ancestral and derived alleles among them (Supplemental Figure S7a). We cannot observe loci in which all chromosomes are identical, leaving 16 patterns that contain one to four derived alleles. Population genetic principles state that derived alleles have systematically lower allele frequencies than ancestral ones. Indeed, for both insertions and deletions, the SV loci that have one derived allele (and four ancestral ones) are much more common than those that have 2 or more (Supplemental Figure S7b). In 12 of the segregation patterns, genotypes of NA12878 and NA24385 differ. These can be arranged in 6 pairs where the individuals' genotypes are equivalent, for example, one heterozygote and one ancestral homozygote, with the reference carrying a derived allele (rightmost green box in Supplemental Figure S7a). Distinguishing insertions and deletions, this gives 12 classes (bottom of Figure 3a), each of which has two SV counts that should be very similar to each other assuming that the two individuals do not come from very different populations. Indeed, the correlation between these equivalence classes is very high ($R^2$=0.92, Supplemental Figure S7c,d).

#### Supplemental Tables and Figures

| Sequenced Library | Sample ID | Raw fold-coverage | $\mu_{FL}$ / W$\mu_{FL}$ (kb) | PCR duplication (%) | $C_F$ (X) |
|---|---|---|---|---|---|
| L1 | NA12878 | 103 | 79.0 / 304.3 | 19.97 | 123 |
| L2 | NA12878 | 192 | 24.0 / 41.1 | 10.92 | 334 |
| L3 | NA12878 | 106 | 99.2 / 214.5 | 11.09 | 958 |
| L7 | NA12878 | 78 | 37.4 / 82.3 | 4.84 | 243 |
| L4 | NA24385 | 100 | 120.8 / 267.4 | 18.51 | 208 |
| L5 | NA24385 | 100 | 64.2 / 151.7 | 12.39 | 803 |
| L6 | NA24385 | 100 | 92.1 / 216.9 | 10.88 | 1504 |

**Supplemental Table S1**. Key parameters of our 10x libraries from NA12878 and NA24385, L1 to L6, and a previously published library from 10x Genomics (L7).

| SNP | | True Positives | False Negatives | False Positives | Genotype Mismatch | Precision | Recall | F1 |
|---|---|---|---|---|---|---|---|---|
| L1 (NA 12878) | Aquila | 2,873,378 | 169,402 | 140,812 | 5,209 | 0.953 | 0.944 | 0.949 |
| | FreeBayes | 3,020,505 | 22,278 | 86,215 | 6,746 | 0.972 | 0.993 | 0.982 |
| | Longranger | 3,038,019 | 4,764 | 294,103 | 2,870 | 0.911 | 0.998 | 0.953 |
| L2 (NA 12878) | Aquila | 2,941,087 | 101,696 | 322,480 | 16,709 | 0.901 | 0.967 | 0.932 |
| | FreeBayes | 3,038,255 | 4,528 | 29,288 | 3,657 | 0.990 | 0.999 | 0.994 |
| | Longranger | 3,028,310 | 14,473 | 111,869 | 2,181 | 0.964 | 0.995 | 0.980 |
| L7 (NA 12878) | Aquila | 2,965,257 | 77,524 | 265,428 | 10,521 | 0.918 | 0.975 | 0.945 |
| | FreeBayes | 3,033,390 | 9,393 | 55,835 | 5,576 | 0.982 | 0.997 | 0.989 |
| L4 (NA 24385) | Aquila | 2,962,316 | 67,042 | 129,018 | 9,291 | 0.958 | 0.978 | 0.968 |
| | FreeBayes | 3,015,793 | 13,565 | 45,759 | 5,507 | 0.985 | 0.996 | 0.990 |
| | Longranger | 3,024,732 | 4,626 | 149,327 | 2,244 | 0.953 | 0.998 | 0.975 |
| L6 (NA 24385) | Aquila | 2,985,796 | 43,561 | 108,248 | 3,952 | 0.965 | 0.986 | 0.975 |
| | FreeBayes | 3,021,607 | 7,751 | 52,638 | 3,846 | 0.983 | 0.997 | 0.990 |
| | Longranger | N/A | | | | | | |

**Supplemental Table S2**. Accuracy of SNP calling, comparing assembly-based calling with two mapping-based approaches on the same libraries' linked read data. The benchmark is GiaB v3.3.2. Variant counts and performance scores were generated by RTGtools/hap.py, an Illumina haplotype comparison/benchmarking tool. Longranger calls were executed with "-vcmode=gatk". N/A, not applicable because Longranger could not complete runs in wgs mode.

| Small Indels | | True Positives | False Negatives | False Positives | Genotype Mismatch | Precision | Recall | F1 |
|---|---|---|---|---|---|---|---|---|
| **NA12878 (hg38)** | **L1** | 475,455 | 55,927 | 67,656 | 27,554 | 0.875 | 0.895 | 0.885 |
| | **L2** | 475,000 | 56,382 | 40,802 | 16,726 | 0.921 | 0.894 | 0.907 |
| | **L3** | 499,301 | 32,081 | 40,292 | 9,493 | 0.925 | 0.940 | 0.932 |
| | **L7** | 483,848 | 47,534 | 66,081 | 20,154 | 0.880 | 0.911 | 0.895 |
| **NA24385 (hg38)** | **L4** | 469,972 | 33,081 | 59,299 | 11,365 | 0.888 | 0.934 | 0.911 |
| | **L5** | 476,139 | 26,914 | 35,315 | 7,986 | 0.931 | 0.947 | 0.939 |
| | **L6** | 474,306 | 28,747 | 36,944 | 8,353 | 0.928 | 0.943 | 0.935 |
| | **L5+L6** | 473,895 | 29,158 | 15,724 | 5,335 | 0.968 | 0.942 | 0.955 |
| **NA24385 (hg19)** | **L5** | 467,492 | 25,108 | 33,250 | 7,607 | 0.934 | 0.949 | 0.941 |
| | **L5** | 465,488 | 27,112 | 34,674 | 8,037 | 0.931 | 0.945 | 0.938 |
| | **L5+L6** | 467,624 | 24,976 | 14,512 | 4,777 | 0.970 | 0.949 | 0.959 |

**Supplemental Table S3**. Accuracy of Aquila small indel calls by benchmarking against the GiaB small indel callset v4.0.

| Library | Homo Aq | Homo Fb | Hetero Aq | Hetero Fb | Total Aq | Total Fb |
|---|---|---|---|---|---|---|
| L1 | 1,437,914 | 1,445,653 | 2,376,367 | 2,508,972 | 3,814,281 | 3,954,625 |
| L2 | 1,429,006 | 1,462,536 | 2,713,879 | 2,486,207 | 4,142,885 | 3,948,743 |
| L3 | 1,460,820 | 1,453,991 | 2,510,624 | 2,495,730 | 3,971,444 | 3,949,721 |
| L7 | 1,433,595 | 1,448,088 | 2,687,428 | 2,492,822 | 4,122,993 | 3,940,910 |
| L4 | 1,405,855 | 1,470,476 | 2,494,627 | 2,465,147 | 3,900,482 | 3,935,623 |
| L5 | 1,409,588 | 1,481,046 | 2,473,281 | 2,480,638 | 3,882,869 | 3,961,684 |
| L6 | 1,403,142 | 1,479,839 | 2,478,552 | 2,482,312 | 3,881,694 | 3,962,151 |

**Supplemental Table S4**. Comparison of the number of SNP calls, in diploid regions, using pairwise contig-to-reference alignment by Aquila (Aq) versus FreeBayes (Fb) calls on the LongRanger-aligned reads. Homo = homozygous, Hetero = heterozygous.

| Library | Deletions | | | Insertions | | |
|---|---|---|---|---|---|---|
| | Homo | Hetero | Total | Homo | Hetero | Total |
| L1 | 116,003 | 387,479 | 503,482 | 115,136 | 374,926 | 490,062 |
| L2 | 110,106 | 383,338 | 493,444 | 109,712 | 356,245 | 465,957 |
| L3 | 131,602 | 385,741 | 517,343 | 130,004 | 359,966 | 489,970 |
| L7 | 115,593 | 414,024 | 529,617 | 113,559 | 366,859 | 480,418 |
| L4 | 125,019 | 394,529 | 519,548 | 124,243 | 379,019 | 503,262 |
| L5 | 127,593 | 383,532 | 511,125 | 126,775 | 357,672 | 484,447 |
| L6 | 125,692 | 382,402 | 508,094 | 125,042 | 355,661 | 480,703 |

**Supplemental Table S5**. Number of assembly-based small indel calls (< 50bp) in diploid regions for each library. Deletions: homozygous ones, heterozygous ones, and total number. Insertions: homozygous ones, heterozygous ones, and total number.

| Deletions | Homo | Hetero | Total | Insertions | Homo | Hetero | Total |
|---|---|---|---|---|---|---|---|
| L1 | 2,427 | 13,953 | 16,380 | L1 | 576 | 16,465 | 17,041 |
| L2 | 3,072 | 13,222 | 16,294 | L2 | 700 | 5,751 | 6,451 |
| L3 | 3,398 | 14,408 | 17,806 | L3 | 772 | 5,285 | 6,057 |
| L7 | 2,849 | 15,922 | 18,771 | L7 | 632 | 9,338 | 9,970 |
| L4 | 2,952 | 14,962 | 17,914 | L4 | 618 | 10,086 | 10,704 |
| L5 | 3,254 | 14,131 | 17,385 | L5 | 761 | 5,295 | 6,056 |
| L6 | 3,294 | 14,594 | 17,888 | L6 | 757 | 5,009 | 5,766 |

**Supplemental Table S6**. Number of SVs >= 50bp in diploid regions for each library. Deletions: homozygous ones, heterozygous ones, and total number. Insertions: homozygous ones, heterozygous ones, and total number.

| | SNPs (Initial phase) | SNPs (Inferred) | Indels (Inferred) | SVs (Inferred) | Total |
|---|---|---|---|---|---|
| **L1** | 1,963,497 | 578,558 | 698,828 | 29,795 | 3,270,678 |
| **L2** | 2,094,973 | 543,558 | 649,789 | 17,123 | 3,305,443 |
| **L3** | 1,746,930 | 802,233 | 674,018 | 18,895 | 3,242,076 |
| **L4** | 1,885,560 | 640,520 | 684,301 | 23,544 | 3,233,925 |
| **L5** | 1,973,507 | 538,224 | 660,832 | 18,373 | 3,190,936 |
| **L6** | 1,715,391 | 801,564 | 656,335 | 18,469 | 3,191,759 |

**Supplemental Table S7**. Number of phased variants for all libraries. Number of heterozygous SNPs phased by the Aquila haplotyping algorithm (SNPs, Initial phase), number of heterozygous SNPs phased by inference (SNPs, Inferred), number of heterozygous small Indels (< 50*bp*) phased by inference (Indels, Inferred), number of heterozygous SVs (>=50*bp*) phased by inference (SVs, Inferred), and the total number of heterozygous variants.

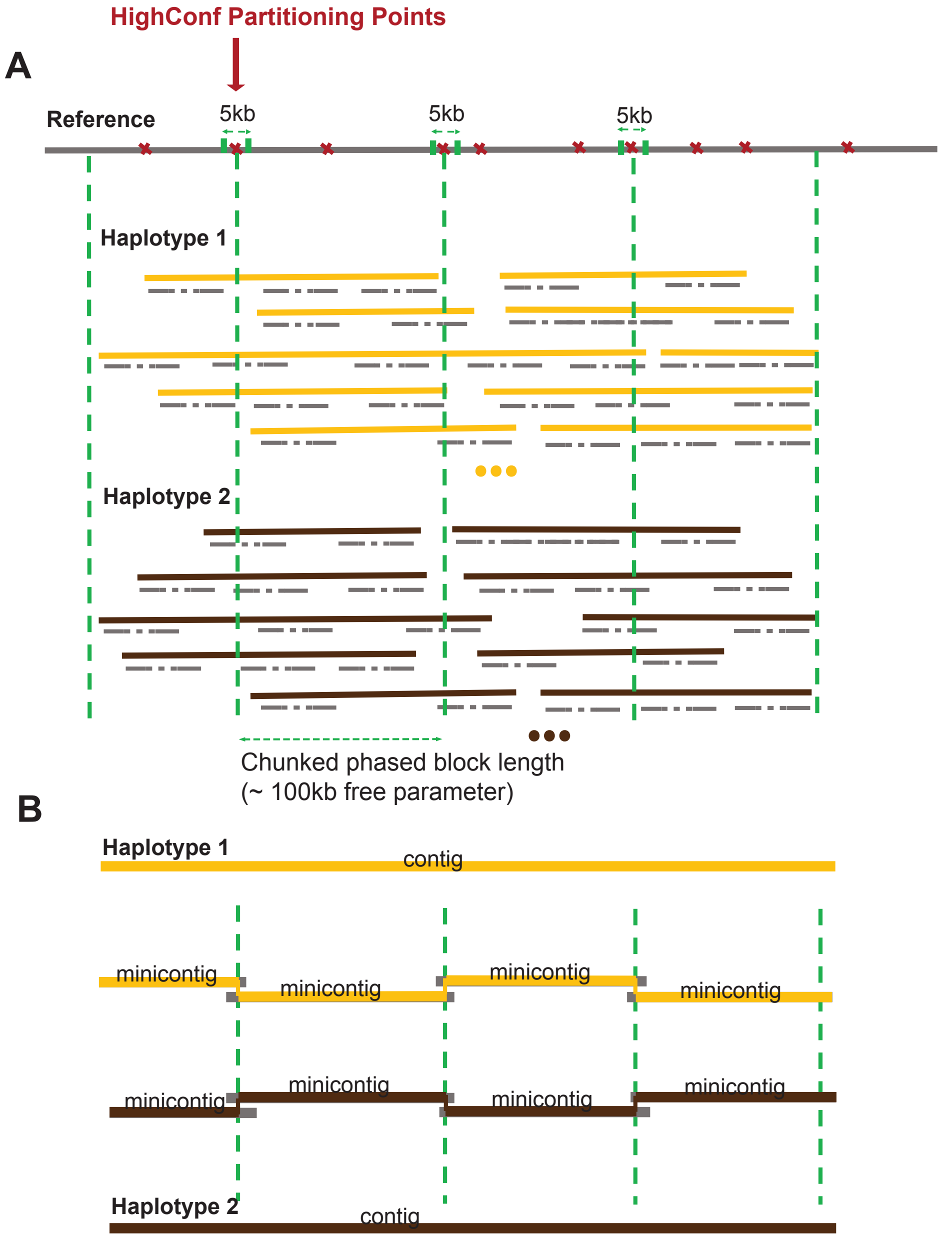


**Supplemental Figure S1**: High confidence partitioning points. A. Cutting one big phase block (in this example, ~ 300kb) from each haplotype into three small chunks to assemble "minicontigs" (in this example, ~100kb each) based on the high-confidence partitioning profile. There is a 5kb shift tolerance to allow Aquila find the high confidence partitioning point. B. Concatenation of minicontigs from each small chunk into final contig for both haplotypes within one big phase block.

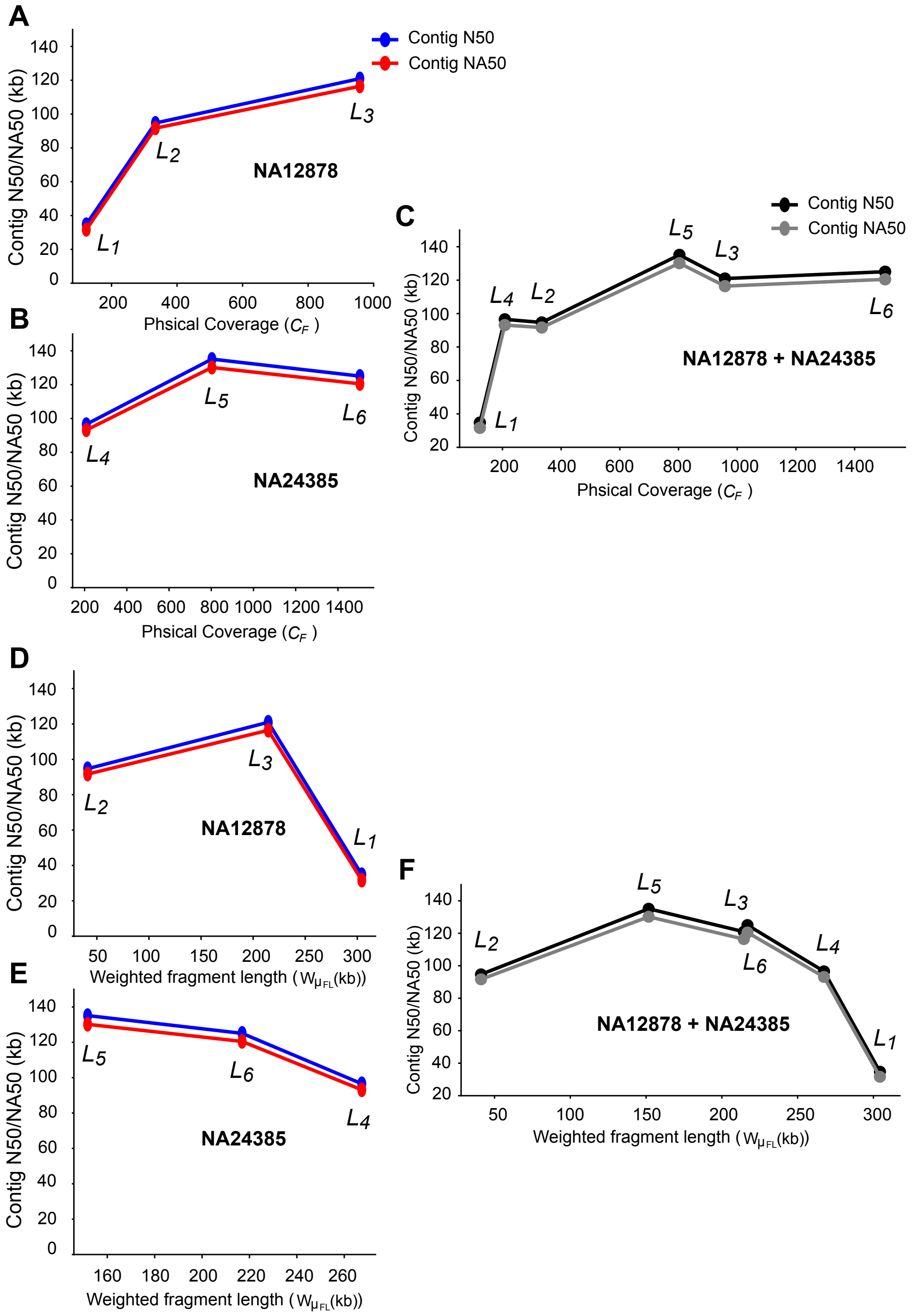


**Supplemental Figure S2**: A, B, C. Contig lengths as a function of physical coverage ($C_F$); D, E, F. Contig lengths as a function of weighted fragment length ($W_{\mu FL}$).

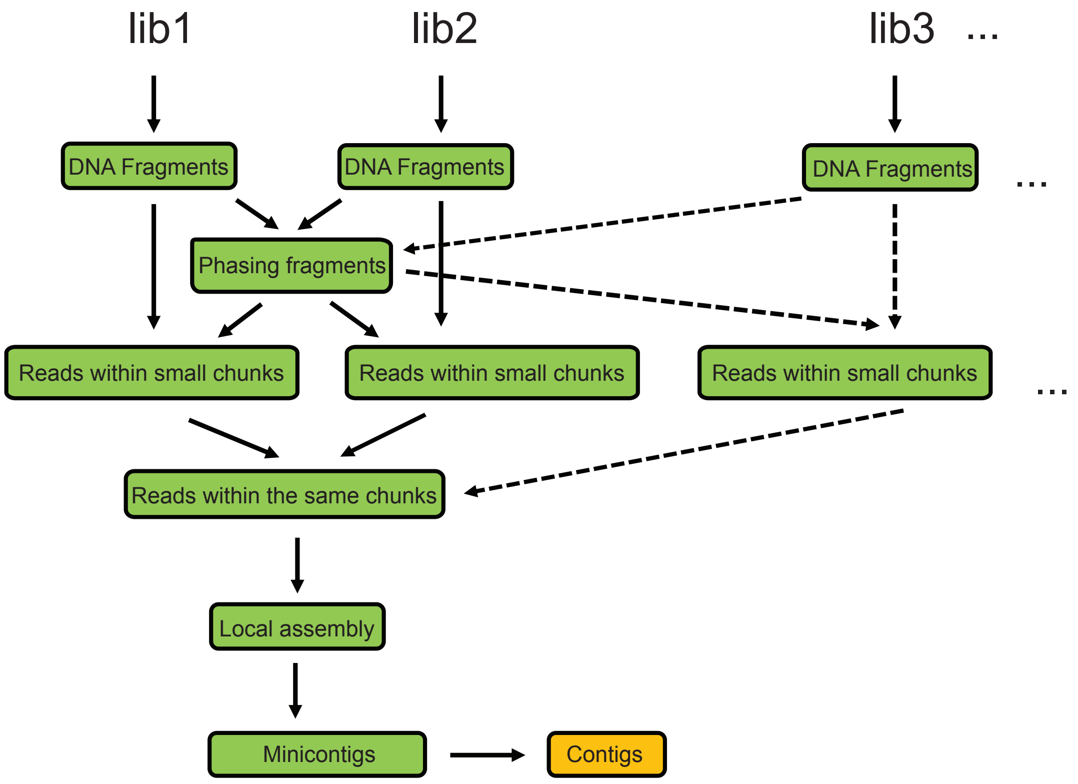


**Supplemental Figure S3**: Assemblies of multiple linked-read libraries. Long DNA molecules/fragments are first reconstructed from each library separately, but then the workflow is exactly the same as the single-library pipeline: The haplotyping module is applied to all the fragments together, short reads are extracted for each library based on the final phased information and then accumulated for the same chunks, followed by local assembly to generate minicontigs. Finally, minicontigs are iteratively concatenated into final contigs.

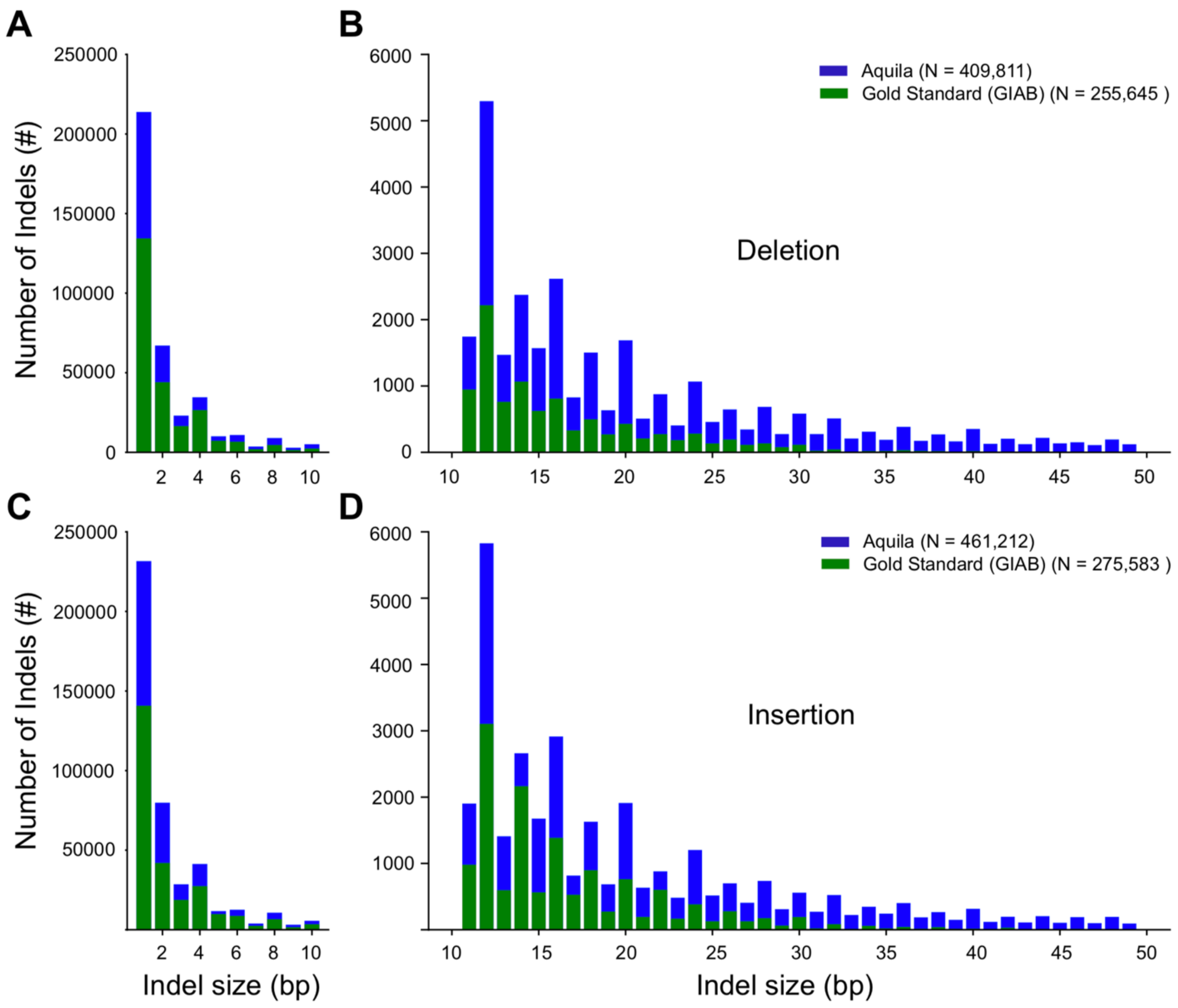


**Supplemental Figure S4**: Small indel size distribution of NA12878 for library 3 (blue plus green), with the distribution for GiaB benchmark in green only for comparison. A, deletions (<= 10bp); B, deletions (> 10bp and < 50bp); C, insertions (<= 10bp); D, insertions (> 10bp and < 50bp).

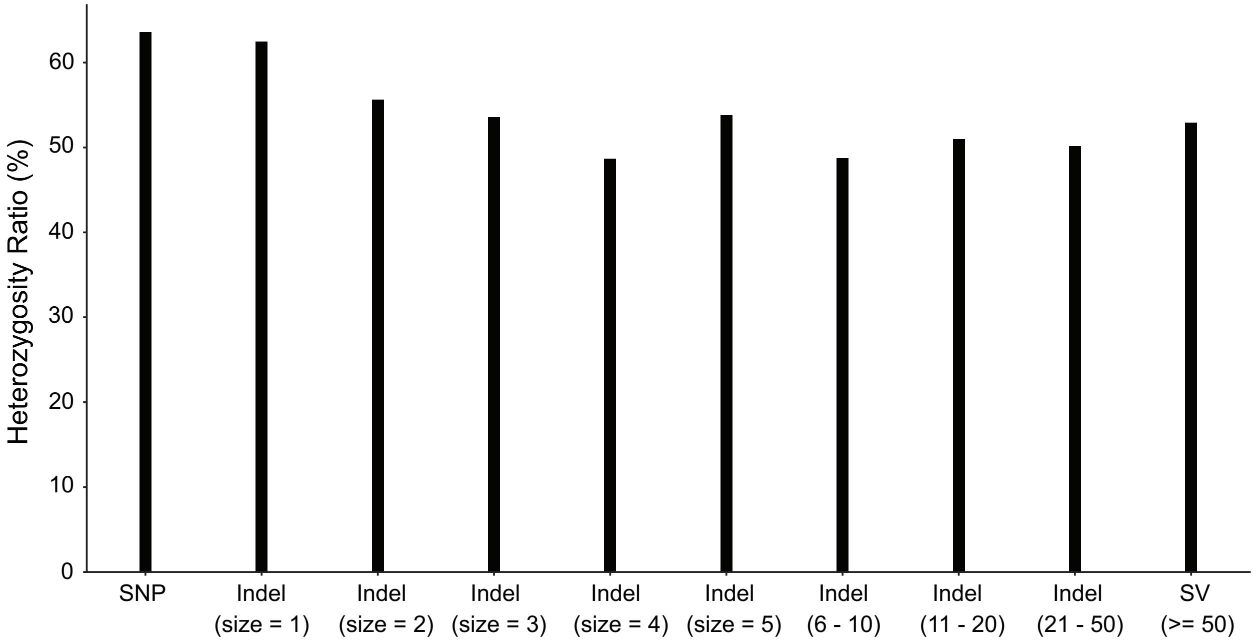


**Supplemental Figure S5**: Average heterozygosity of all types and sizes of discovered variants.

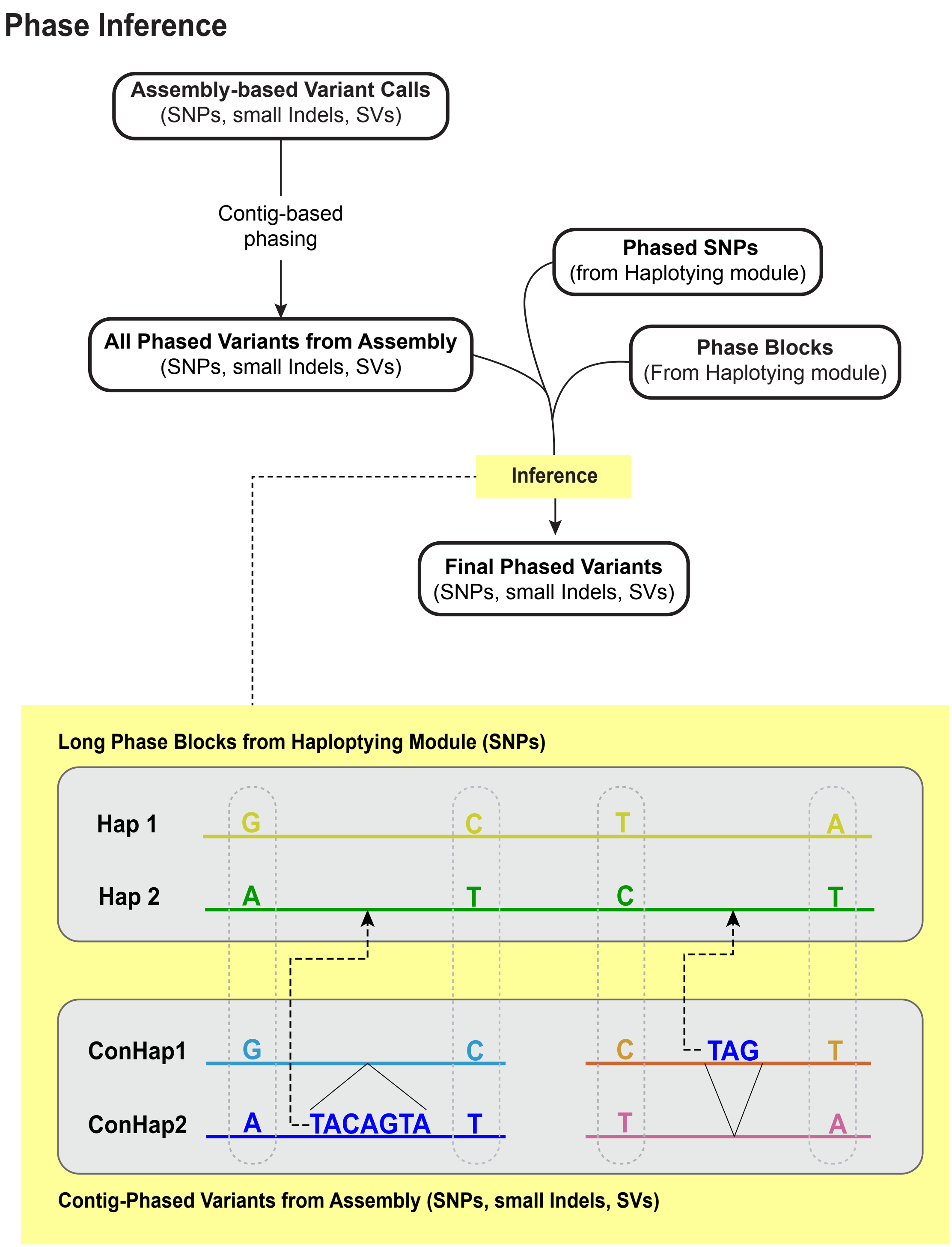


**Supplemental Figure S6**: Inference of phase for assembly-discovered variants on the basis of phase blocks generated in the Haplotyping module.

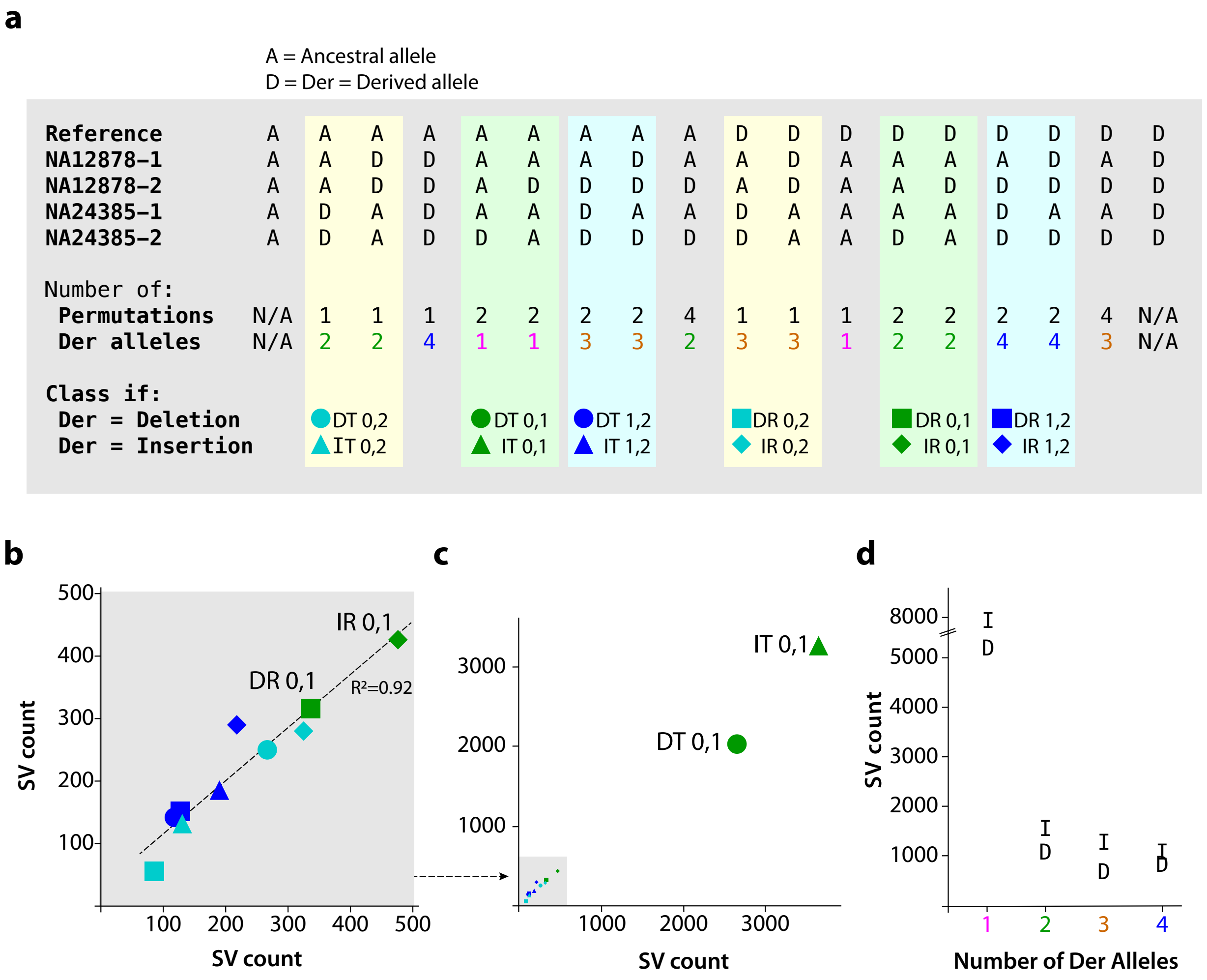


**Supplemental Figure S7**. Frequencies and genetic configurations of detected actual insertions and deletions as determined by alignment to ape genomes. (a) Each column depicts a possible configuration of ancestral and derived alleles in the five haplomes; all possibilities are shown but we cannot detect variation shown in the first and last columns as all sequences are identical in those cases. Each pattern is distinct but, due to the nature of sampling, represents between 1 and 4 possible arrangements. The number of derived alleles in each pattern is also indicated. Shorthand for symbols indicating the molecular type and who carries the derived alleles (MD) are: DT, deletion carried by a target; DR, deletion carried by the reference; IT, insertion carried by a target, IR, insertion carried by the reference. 0,1: single heterozygote; 0,2: single homozygote; 1,2: one target is a heterozygote, the other a homozygote. (b,c) MD numbers in the individuals correlated to each other. Symbols represent classes as described in (a). For example, the green diamond (IR 0,1) indicates SVs defined by the fact that the derived allele is an insertion ("I") in the reference sequence ("R"), designated as "IR". There are 475 SVs in which NA12878 is homozygous for the reference allele and NA24385 is heterozygous for the derived allele, and 428 SVs for which NA12878 is heterozygous for the derived allele and NA24385 is homozygous for the reference allele. All other data points are analogous, always comparing NA12878 calls to their independent and equivalent counterparts.
